# An automated pipeline for extracting histological stain area fraction for voxelwise quantitative MRI-histology comparisons

**DOI:** 10.1101/2022.02.10.479718

**Authors:** Daniel Z.L. Kor, Saad Jbabdi, Istvan N. Huszar, Jeroen Mollink, Benjamin C. Tendler, Sean Foxley, Chaoyue Wang, Connor Scott, Adele Smart, Olaf Ansorge, Menuka Pallebage-Gamarallage, Karla L. Miller, Amy F.D. Howard

**Author notes:** Address for Correspondence: Daniel Zhi Liang Kor, Wellcome Centre for Integrative Neuroimaging, FMRIB John Radcliffe Hospital, Headington, Oxford OX3 9DU, UK. Contributed equally (Karla L. Miller, Amy F.D. Howard).

## Abstract

The acquisition of MRI and histology in the same post-mortem tissue sample enables direct correlation between MRI and histologically-derived parameters. However, there still lacks a standardised automated pipeline to process histology data, with most studies relying on heavy manual intervention. Here, we introduce an automated pipeline to extract a quantitative histological measure for staining density (stain area fraction) from multiple immunohistochemical (IHC) stains. The pipeline is designed to directly address key IHC artefacts related to tissue staining and slide digitisation. Here, the pipeline was applied to post-mortem human brain data from multiple subjects, relating MRI parameters (FA, MD, R2*, R1) to IHC slides stained for myelin, neurofilaments, microglia and activated microglia. Utilising high-quality MRI-histology coregistrations, we then performed whole-slide voxelwise comparisons (simple correlations, partial correlations and multiple regression analyses) between multimodal MRI- and IHC-derived parameters. The pipeline was found to be reproducible, robust to artefacts and generalisable across multiple IHC stains. Our partial correlation results suggest that some simple MRI-SAF correlations should be interpreted with caution, due to the co-localisation of certain tissue features (e.g. myelin and neurofilaments). Further, we find activated microglia to consistently be the strongest predictor of DTI FA, which may suggest sensitivity of diffusion MRI to neuroinflammation. Taken together, these results show the utility of this approach in carefully curating IHC data and performing multimodal analyses to better understand microstructural relationships with MRI.

## 1 Introduction

Magnetic resonance imaging (MRI) is a powerful tool that can be used to evaluate neurodegenerative disorders in-vivo. MRI techniques have produced quantitative parameters sensitive to macroscopic neuropathological changes [1-3]. However, MRI parameters are generally non-specific and sensitive to multiple factors related to tissue microstructure. Coupled with millimetre resolution, this leads to difficulty in determining the microstructural underpinnings of a given MRI change (e.g., in disease).

Immunohistochemistry (IHC) is a histological staining technique that can be used to address this difficulty. IHC uses primary antibodies to stain target antigens (proteins related to microstructural features of interest) with high specificity. The distributions of antigens are then commonly visualised using the chromogen 3’3-diaminobenzidine (DAB) to stain the marked proteins brown. To add contextual information and localise stained tissue features, haematoxylin is often used as a counterstain to mark cell nuclei purple. The acquisition of IHC and MRI in the same post-mortem tissue sample enables direct correlation of MR and histologically-derived metrics. This is a common methodology used to validate MR parameters and improve our microstructural interpretation of MRI. Here, post-mortem MRI functions as a crucial intermediary between the IHC data and in-vivo imaging. Post-mortem MRI shares a common tissue state with IHC, while possessing the same signal forming mechanisms with in-vivo MRI [4].

While many studies relate IHC to MR parameters [5-13], there is still a lack of automated pipelines for extracting quantitative metrics from IHC slides of neuronal tissue [14]. Most MRI-IHC analyses rely on heavy manual intervention [14-17], with some using semi-quantitative metrics, such as staining intensity scores resembling *low, moderate* or *strong* [18,19]. More quantitative metrics can be derived via “colour deconvolution” to provide a more detailed description of IHC staining. When multiple stains have been applied to the same tissue section, colour deconvolution utilises the stains’ colour information to separate the densities for different stain channels (e.g. separating the DAB-stained densities from background haematoxylin) [20]. Some IHC analyses use the DAB channel’s density to approximate the amount of targeted protein within the tissue [18, 21-23]. However, this interpretation is limited as the densities do not scale linearly with protein concentration [17, 22, 24]. To circumvent this, other pipelines extract the stain area fraction (SAF), i.e. the number of DAB-stained pixels within a given area. In most pipelines, SAF is quantified by manually setting a threshold for the DAB channel to segment microstructural tissue compartments from non-specific background staining in regions-of-interest (ROIs) [14-17].

The manual derivation of SAF has two main issues. First, the manually-set threshold may not be optimal for any given slide. Manual thresholds are dependent on the operator’s expertise. During this process, a single threshold is often applied to all slides in a dataset or a batch of slides which have been processed together [9]. Although this is time efficient and avoids intra-observer variability, it comes at the expense of optimising thresholds for individual slides, resulting in decreased robustness to histological artefacts, such as slide-to-slide and within-slide staining intensity variation. Even when a single tissue preparation protocol is followed, slide-to-slide staining intensity variation is introduced due to variations in tissue sample preparation and staining protocol [25-27], which produce artificial (non-biological) differences in stain intensity and colour information. Examples of within-slide staining artefacts include a gradual staining gradient, with stronger staining at one end of the slide progressing to weaker staining at the other, where the axis of the gradient is related to how the slide is positioned during staining, and striping artefacts from slide digitisation. The latter describes sharp vertical bands of intensity variation or striping across whole slide images, which arise from imperfect illumination and/or the stitching together of multiple strips of the image when scanning the slide [28]. When unaccounted for, these artefacts may impact the extracted IHC metrics.

Second, these manual workflows are time intensive. This restricts research studies to smaller sample sizes (i.e. less slides and/or subjects), and limits IHC analyses to hand-drawn ROIs, as opposed to analysing voxels from the whole slide. Ideally, an automated workflow will allow MRI-SAF correlations to be derived at the MRI voxel level. To achieve this, the workflow will need to be able to produce whole-slide SAF maps from IHC data. These SAF maps can then be co-registered MRI to enable voxelwise analysis.

Here, we propose an automated SAF pipeline, in the context of an end-to-end MRI-histology workflow, to address these challenges. The automated IHC processing pipeline is able to extract SAF maps from IHC-stained slides for myelin, neurofilaments and microglia. The pipeline was first evaluated on an IHC dataset designed to test the pipeline’s reliability. Once evaluated, we applied it on a second dataset containing both IHC and MRI data. We register and correlate these SAF maps on a voxelwise basis with diffusion-MRI (fractional anisotropy, FA; mean diffusivity, MD) and relaxometry (R2*, R1) maps. To account for covariance between stains, we use partial correlation to identify the unique variance in MRI parameters explained by each targeted protein. Finally, we perform multiple regression with all stains to derive a predictive model of each MR parameter, which may be driven by multiple microstructural sources.

## 2 Data Acquisition

### 2.1 Immunohistochemistry data

We applied the SAF pipeline to two datasets: one that was acquired specifically to evaluate the reliability of our pipeline and which includes only IHC data, and a second, previously described dataset that includes both IHC and co-registered MRI [9]. Both datasets contain tissue from the same 15 post-mortem brains of patients diagnosed with amyotrophic lateral sclerosis (ALS) and healthy controls (CTL) (12 x ALS, 3 x CTL). All antibodies were visualised with DAB and the slides were counterstained with hematoxylin. Slides were digitised with the Aperio ScanScope® AT Turbo (Leica Biosystems) at x20 object magnification (0.5 µm/pixel).

#### 2.1.1 Evaluation dataset

The “evaluation dataset” was acquired to evaluate our pipeline’s performance in terms of reproducibility and robustness to key histological artefacts. Ideally, this data can distinguish true biological variation (e.g. due to differences in pathology, which is assumed to be consistent between adjacent IHC slides) from variance related to staining artefacts and analysis. Consequently, data was collected from adjacent tissue slides. Our analyses using the evaluation dataset assume that these slides have similar tissue microstructure and that the true biological between-slide variance is low.

For each of the 15 brains, 12-15 consecutive slides (6-μm thick) were obtained from the primary motor cortex (face region). These adjacent slides were separated into groups of 4-5 slides. Each group was stained for either activated microglia/macrophages (CD68), myelin (PLP) or phosphorylated neurofilaments (SMI312). The staining protocol is identical to that in [9] and full staining protocols are provided in Appendix A. Stains were done in batches to further reduce technical variation. For the evaluation of reproducibility, these slides were first manually quality-checked on a per slide basis. The criterion was based on whether the slides contained artefacts that will not yield useful SAF estimates from both manual and automated pipeline. These include excessively illuminated slides, significant amounts of inconsistent staining, and slides that have been sectioned poorly.

#### 2.1.2 Multiple-region dataset

We applied our SAF pipeline (Section 3.1) on the “multiple-region dataset” [9] to generate SAF maps. This multiple-region dataset includes IHC and co-registered MRI data from multiple brain regions with varying levels of disease pathology. Both the multiple-region dataset and its SAF maps will be made available in a future release of the Digital Brain Bank [4]. We consider data from the visual cortex (both hemispheres), anterior cingulate (cingulum bundle, corpus callosum) and hippocampus (HC). Slides were stained to visualise CD68, Iba1 (all microglia), PLP and SMI312. Prior to any analysis, all slides were quality-checked similarly to the evaluation dataset.

### 2.2 MRI data

The data acquisition and preprocessing procedures of post-mortem MRI data have been previously described ([9], [11], [29]). In brief, whole brains were imaged in a 7T human scanner (Siemens Healthcare, Erlangen, Germany) using a 1Tx/32Rx head coil. R2* maps were estimated from susceptibility-weighted data acquired with a 3D multi-echo GRE sequence (parameters: TEs=2, 8.6, 15.2, 21.8, 28.4, 35 ms with monopolar readout and non-selective RF pulse, TR=38 ms, flip angle=15°, bandwidth=650 Hz/pixel, and in-plane resolution=0.5 × 0.5 mm^2^ [11]). R1 maps were estimated from T1-weighted data collected with a multi-TI turbo spin-echo protocol (typical parameters: TE=14.0 ms, TR=1000 ms, TIs=60, 120, 240, 480, 935 ms, flip angles=90° and 180°, bandwidth=130 Hz/pixel, and in-plane resolution=1.0 × 1.0 mm^2^ [9]). Diffusion tensor maps including FA and MD [30] were estimated from a diffusion-weighted steady-state free precession sequence (parameters: TE=21.0 ms, TR=28.0 ms, flip angles=24° and 94°, bandwidth=393 Hz/pixel, q value=300 cm^-1^, number of directions/flip angle=120, and in-plane resolution=0.85 × 0.85 mm^2^ [29]).

### 2.3 Data co-registration

MRI and histology data were previously co-registered using FSL’s Tensor Image Registration Library (TIRL), which is a general-purpose image registration framework designed for MRI-microscopy coregistration [31]. Here, we used TIRL to 1) register PLP with structural MRI (2D-3D) and 2) co-register other stains to PLP (2D-2D). PLP was chosen as the reference microscopy data due to its strong white/grey matter (WM/GM) contrast. All MR parameters maps were first aligned to structural MRI using FLIRT [38]. The generated warps were combined with TIRL to map these MR parameters to the IHC SAF maps for voxelwise correlations.

## 3 Methods

Section 3.1 first provides an overview of the end-to-end workflow for MRI-SAF comparisons. At the centre of this workflow is our automated SAF pipeline, which is described in Section 3.2. The pipeline was designed to be generalisable to multiple IHC stains, reproducible and robust to common IHC artefacts, which we evaluate in Section 3.2.4. The pipeline was then applied to the multiple-region dataset which includes IHC and co-registered MRI, facilitating voxelwise MRI-SAF comparisons. We describe how we evaluated the quality of the co-registration (Section 3.3), and performed voxelwise MRI-SAF correlation and regression analysis (Sections 3.4 & 3.5) to disentangle the contributions of multiple microstructural features stained with IHC to each MR parameter.

### 3.1 The MRI-SAF workflow

Figure 1 shows an overview of the end-to-end MRI-SAF workflow. The SAF pipeline receives high-resolution (0.5 µm/pixel) IHC slides and outputs SAF maps at a target resolution. Utilising precise MRI-SAF co-registrations generated by the Tensor Image Registration Library (TIRL) [31], we perform MRI-SAF analyses to investigate relationships between multiple MR parameters and multiple proteins at a voxelwise level.

**Figure 1:**
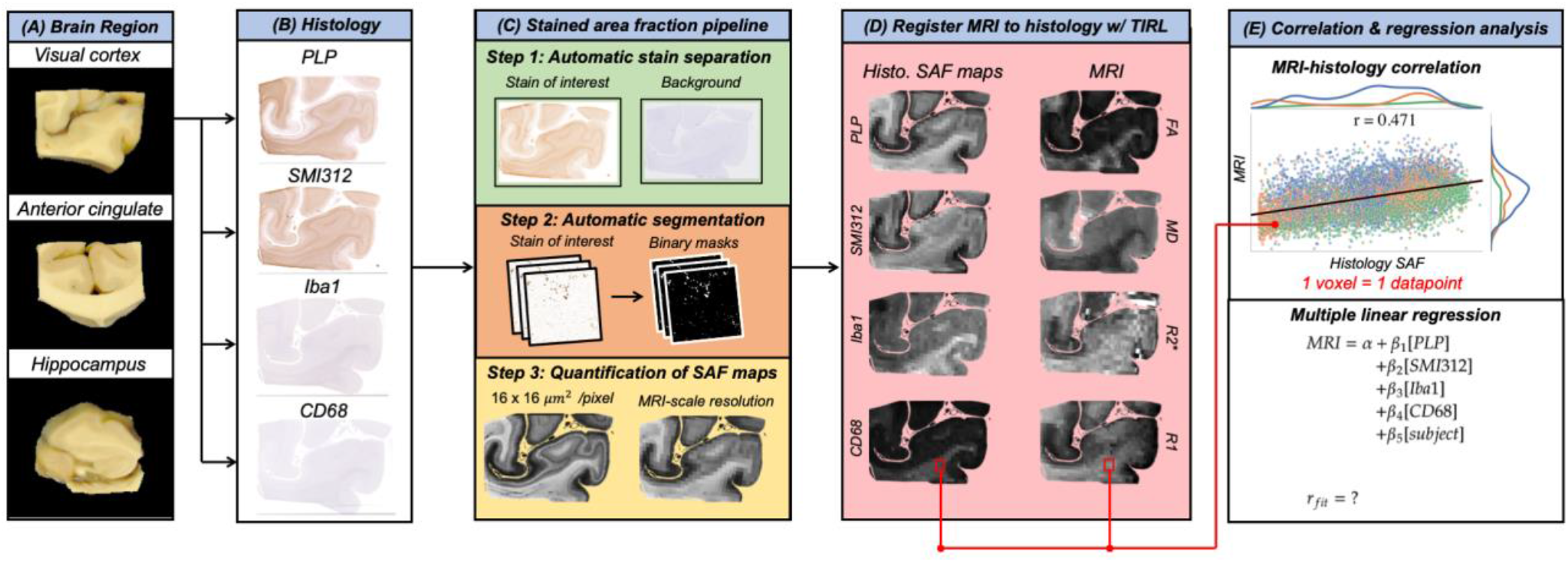
An overview of the MRI-SAF workflow for multiple-region assessment. A: Brain regions are sectioned for immunohistochemistry (IHC). B: Slides are stained for myelin (PLP), neurofilament (SMI312), activated microglia (CD68) & all microglia (Iba1). C: IHC slides are processed to output stain area fraction (SAF) maps (described in section 3.2). D: SAF & MRI maps are then co-registered voxelwise via TIRL. E: Voxels are pooled to perform pairwise correlations and multiple regression between MRI & SAF. Each datapoint in the scatter plot represents a single voxel; the colours represent data from different brain regions (green for visual cortex, orange for anterior cingulate and corpus callosum, blue for hippocampus).

### 3.2 Stain Area Fraction Pipeline

The SAF pipeline maps the RGB intensity values of IHC slides to SAF maps. This pipeline is comprised of three steps: 1) separation of the DAB and haematoxylin stains using colour deconvolution, 2) segmentation of the DAB-stained proteins-of-interest from non-specific DAB staining using an intensity threshold, and 3) calculation of the SAF map at a specified resolution. Here, we present SAF maps both at high resolution (16 µm/pixel) and MRI-scale resolution (0.5 mm/pixel). The SAF pipeline is automated and data-driven, enabling rapid analysis of many IHC slides across multiple subjects, regions and stains.

Ideally, the pipeline should be generalisable to all IHC slides stained with DAB. In practice, we found considerable impact of IHC artefacts in some slides. This motivated the development of two ‘configurations’ of the pipeline: the ‘default’ and ‘artefact’ configuration. The ‘default’ configuration (Figure 2) is designed to emulate an expert histologist when deriving SAF maps of IHC slides with no prominent staining gradient and/or striping artefacts. In cases where these artefacts are present, we propose the ‘artefact’ configuration (Figure 3), which automatically adjusts the local thresholds within-slide to account for the impact of these artefacts. This improves upon a more manual approach, where the adjustment of local thresholds would be extremely time-consuming. Both configurations are automated, and differ based on whether local or whole-slide methods are used for Steps (1) and (2). We now describe each step of the IHC processing pipeline in detail.

**Figure 2:**
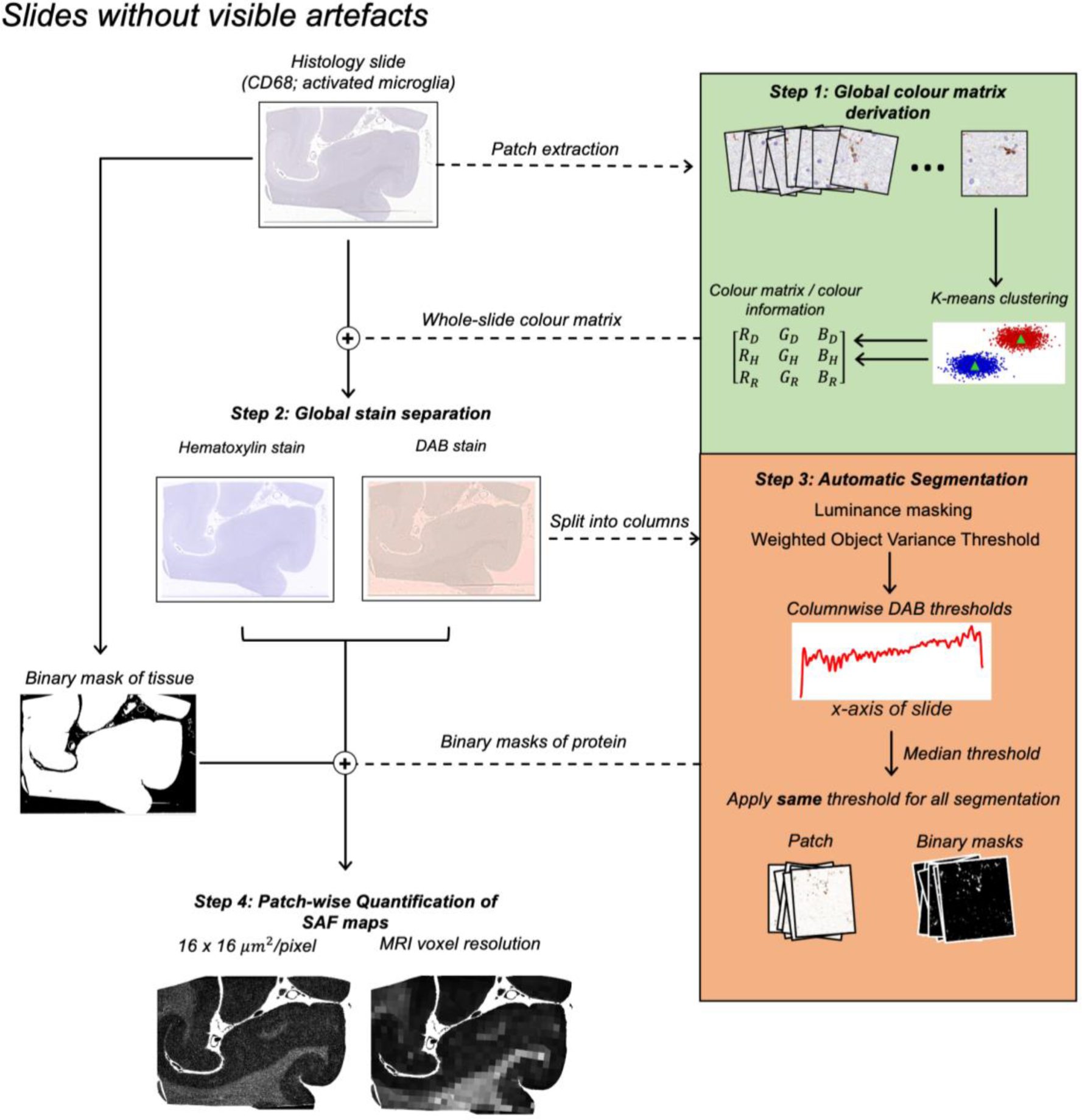
The automated SAF pipeline for stains with little or no intensity gradient and/or stitching artefacts (example: visual cortex). This usually includes slides of stains specifying structures that sparsely populate the brain tissue, such as microglia (Iba1, CD68) and neurofilament (SMI312) in some cases. For each slide, the pipeline 1) derives a single, global colour matrix to separate DAB from haematoxylin, 2) performs stain separation to isolate the DAB channel, 3) automatically segments the DAB channel with a single median threshold, and 4) calculates the SAF at variable resolution.

**Figure 3:**
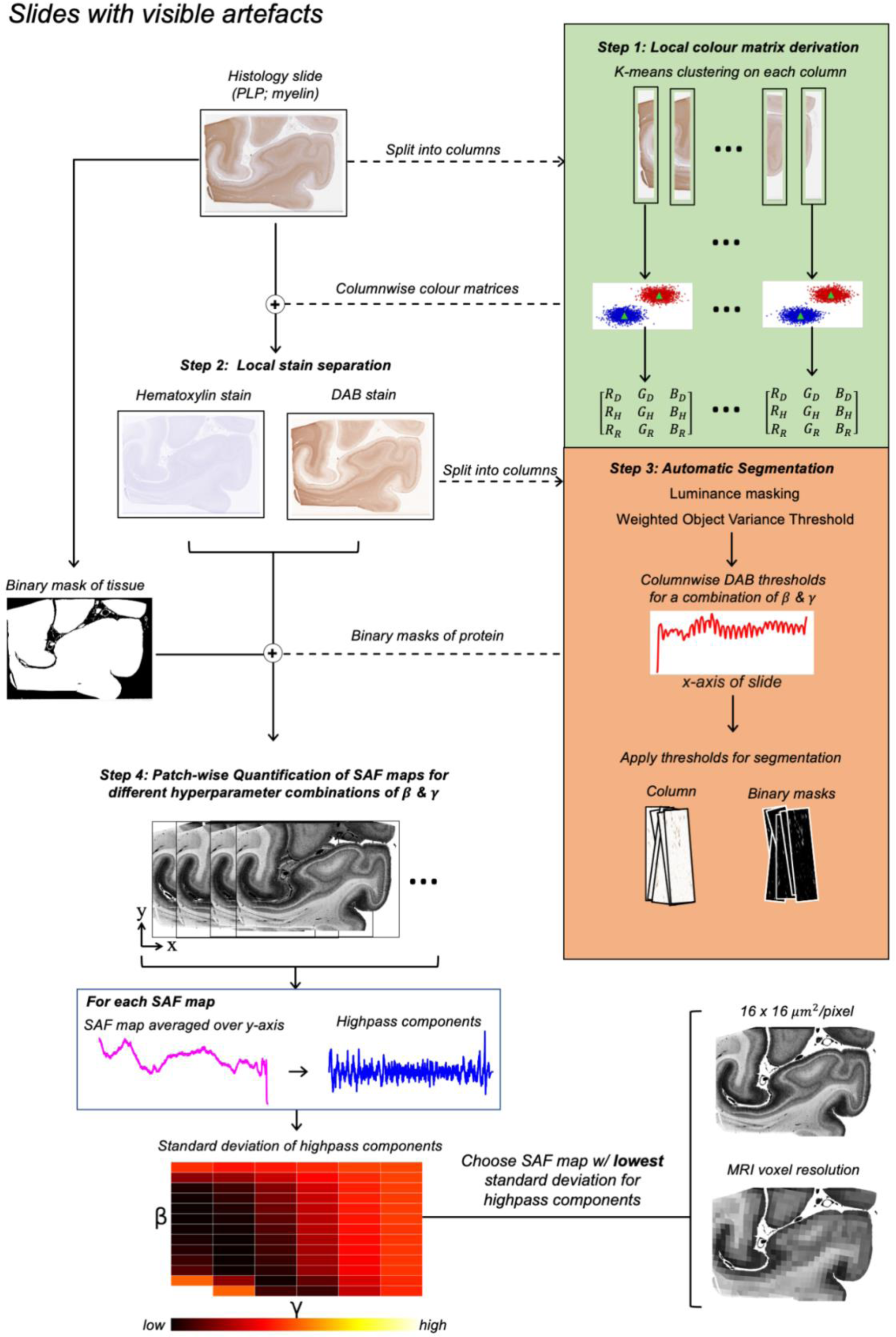
The automated SAF pipeline for stains confounded by staining gradient and/or striping artefacts (visual cortex). For each slide, the pipeline 1) derives multiple, columnar colour matrices from the data, 2) performs stain separation for each column, 3) automatically segments each column’s DAB channel and 4) forms an SAF map. Steps 3 and 4 are repeated for a range of hyperparameters **β** and **γ**, which are optimised via grid-search to account for within-slide artefacts (Section 3.1.2). For more details, please refer to Section 3.2.2. For each SAF map, we averaged the map over its height (y-axis) and chose the SAF map with the lowest standard deviation (i.e. least impacted by these artefacts).

#### 3.2.1 Colour Matrix Derivation for Stain Separation

Colourimetric analysis is based on colour deconvolution, a stain separation method using Beer-Lambert’s law [20]. In Beer-Lambert’s law, the light absorbance (**A**) for RGB channels is linearly related to the density of stain (**C**) via the attenuation coefficients (**ε**). The stain (i.e. DAB, hematoxylin) density in a slide can be computed with a matrix inversion:

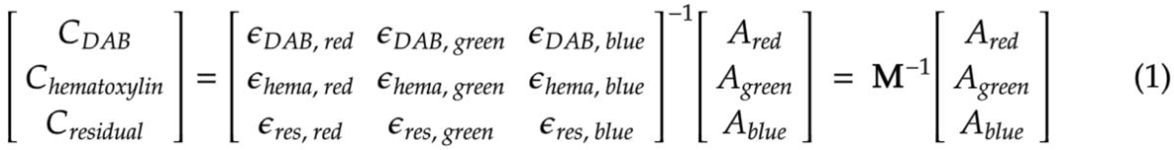

where **M**, termed the colour matrix, is a 3 × 3 matrix with rows defining the stains of interest (**i**), and columns denoting each colour’s attenuation coefficient value (**ε**_**i**,**λ**_). The row is the stain’s colour vector. For more information on Beer-Lambert’s law, see Appendix B.

**M** can be defined from literature [20, 32] or empirically measured from single-stained slides [20]. While it is common to use literature values, using an **M** that is non-specific to each slide leads to poor stain separation and consequently, inconsistent interpretation during slide-to-slide comparison [25, 27, 33]. We address this problem by employing a k-means clustering approach to derive colour information directly from the IHC data. This approach is tailored according to the different configurations.

The ‘default’ configuration is applied to slides without a notable staining gradient. Hence, a single colour matrix, **M**, is sufficient for stain separation in the entire slide. To derive a slide-specific **M**, we used the *k*-means clustering strategy introduced in [34]. In brief, we randomly sampled patches (0.5 × 0.5 mm^2^; n=200) from the slide and performed *k*-means clustering (k=2) to produce 2 cluster centroids corresponding to the colour vectors (i.e. rows in **M**) of hematoxylin and DAB. We then performed a second *k*-means clustering (k=2), this time on the 200 colour vectors output from the first *k*-means. This produced two centroids corresponding to the slide-specific DAB and haematoxylin colour vectors of **M**. Stain separation was then performed using this single M across the whole slide.

In slides with a staining gradient, the **M** artificially changes from one end of the slide to the other. This means a locally changing **M** is required to account for colour differences, as to produce accurate stain separation.

In the ‘artefact’ configuration, we address these staining gradients by applying *k*-means locally along the gradient direction. In our datasets, this staining gradient was present along the horizontal axis of the slide. Hence, we applied *k*-means to columns of data (32-pixels width; height matching the slide) to define **M**_**column**_ on a column-wise basis. Stain separation was performed locally using **M**_**column**_.

For stain separation, an approach emulating NNLS (non-negative least squares), termed the pseudo-NNLS method (see Appendix B), was used to estimate the DAB channel’s stain density (**C**_**DAB**_) according to Equation 1. **C**_**DAB**_ was then converted to intensity values (**I**) using *I* = *log*^*−1*^ _10_ (*C*_*DAB*_).

#### 3.2.2 Automatic Thresholding for Protein Segmentation

The next step is to segment the DAB-stained protein-of-interest from non-specific DAB. Manual analyses require an expert to manually set a threshold. An aim of our pipeline is to derive a data-driven threshold. A common image segmentation approach is Otsu’s method [35], which derives a threshold that maximises the image histogram’s inter-class variance. Our algorithm is based on the related weighted object variance (WOV) algorithm [36], which includes a tunable parameter to separate classes of unequal count and variance. Here, we computed a threshold **t** by minimising over a cost function similar to the one used in WOV:

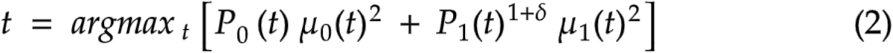

Here, **P**_**i**_**(t)** and **μ**_**i**_**(t)** are the cumulative probabilities and means of the uneven classes i=0 (protein-of-interest) and i=1 (DAB-stained background). As the protein-of-interest may either be the larger (PLP) or smaller (CD68, Iba1) class, a different δ was used for each stain.

To achieve local thresholding, the algorithm was performed on contiguous patches rather than the entire image. Our slides possesed left-to-right gradients in stain density, with vertical bands of intensity (striping). Thresholds **t**_**i**_ were thus calculated on a column-wise basis (32-pixels width; height matching the slide), with **i** representing the column index. A single **δ** was set for each stain, where −1<**δ**<1. When **δ=**0, we obtain the cost function used in Otsu’s method [36], which works optimally when both classes have equal count and variance. By adjusting **δ** to be <0, we were able to accurately segment a protein-of-interest (class 0) with a higher count (PLP) by moving **t** closer to the mean of the DAB-stained background (class 1). Conversely, a **δ**>0 produced a desired threshold for proteins-of-interest with a lower count (CD68, Iba1) by shifting **t** toward the mean of class 0. Before calculating the threshold, the DAB channel is masked to remove the brightest pixels in the column (luminance<0.75) to re-balance the two classes in the DAB channel histogram.

We first describe the ‘default’ configuration of our algorithm. This configuration is applied to slides with no visible striping artefacts; consequently, we concluded that a single threshold is sufficient to segment the entire slide. This fixed threshold is the median threshold of all local WOV thresholds computed column-wise. This data-driven method replaces expert determination of the threshold, **t**, with determination of the hyperparameter, **δ**. Ideally, this hyperparameter is set once by an expert for a stain and will then enable automated, adaptive thresholding on new slides. For our pipeline, we sampled 8-10 patches (0.5 × 0.5 mm^2^) spanning different brain regions, tissue types and subjects to choose a **δ** (per stain) that produced optimal segmentation. Segmentations were vetted with an expert histologist (MPG).

The ‘default’ configuration applies a single threshold aiming to emulate segmentation performed by an expert histologist. However, in slides with variable staining intensity and/or artefacts, a single whole-slide threshold is insufficient for optimal segmentation, even if performed by an expert. We now describe how our ‘artefact’ configuration modifies the ‘default’ configuration to adapt the segmentation threshold to artefacts in a data-driven way.

In slides with striping artefacts, columns affected by striping are characterised by DAB histograms with decreased median absolute deviation (MAD) compared to columns less affected by striping artefacts. Using our algorithm on these narrow histograms results in a threshold that oversegements, artificially inflating the SAF in these columns.

To address this, our pipeline’s ‘artefact’ configuration uses MAD in each column to weight the **δ** to account for how much stitching artefact is present:

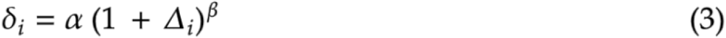

where:

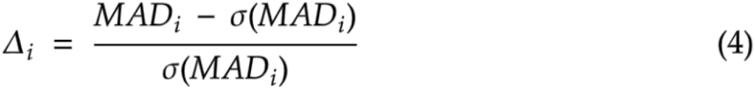

Here, **α** is the main weight attached to the cost function. **MAD**_**i**_ is the median absolute deviation of a DAB channel histogram extracted from the **i**^th^ column. The MAD values are smoothed column-wise with a 1D Gaussian filter (**σ**=16) to produce **σ(MAD**_**i**_), the smoothed MAD in the **i**^th^ column. Hence, **Δ**_**i**_ represents the **i**^th^ column’s change in structure (MAD) relative to neighbouring columns. A large **Δ**_**i**_ represents a sudden, large change in the histogram structure, indicative of the **i**^th^ column being in a striping region. Depending on how much stitching artefact is present, we modulate this weight with a power term **β (**higher **β** used for more prominent striping).

We calculate a column-wise **δ**_**i**_, which is then smoothed using a 1D Gaussian filter (kernel size **γ**) to prevent the thresholds varying abruptly column-to-column. With a given combination of **α, β** and **γ**, we can generate a 1D threshold profile to be applied to the slide’s DAB channel to obtain the segmentation mask. This increase in complexity (from one to three hyperparameters) enables the pipeline to handle slides that cannot be accurately segmented with a single threshold. An example of how these hyperparameters affect the SAF map is shown in Figures S1 and S2 (Appendix C).

The hyperparameter **α** was optimised using the same expert procedure for **δ** (‘default’ configuration), with one value chosen per stain. The hyperparameters **β, γ** were selected from a grid search to minimise the standard deviation of highpass components in the resulting SAF (see Section 3.2.4), and optimised on a per-slide basis.

In our multiple-region dataset, we observed stitching artefacts and staining gradients in the PLP slides only. Hence, we applied the ‘artefact’ configuration on our PLP slides (**α**=-0.60) and the ‘default’ configuration on our CD68 (**δ**=0.05), Iba1 (**δ**=0.05) and SMI312 (**δ**=-0.30) slides. Figure 4 shows some example segmentations.

**Figure 4:**
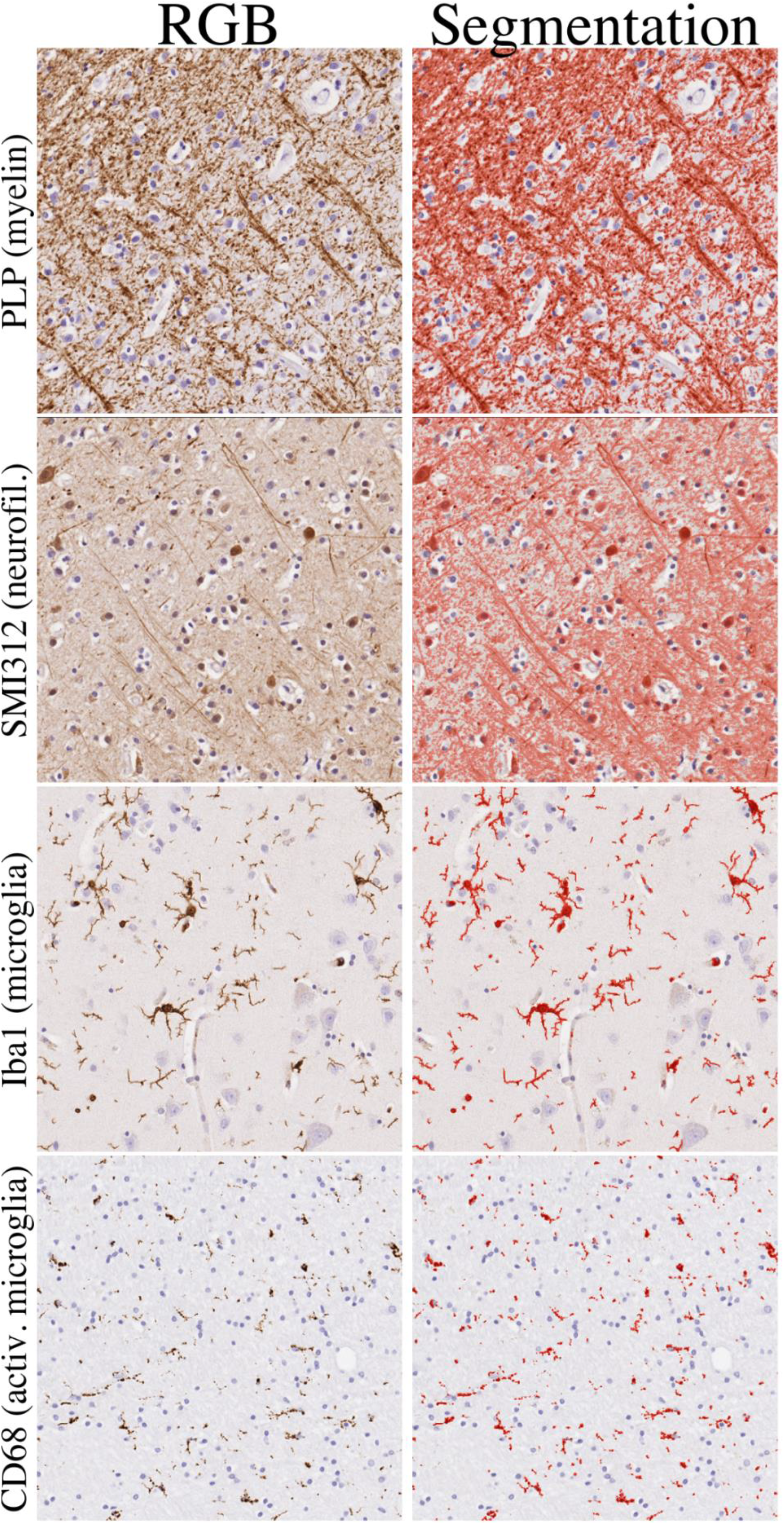
Examples of the segmentation (right column; red overlay) produced from the automated SAF pipeline with optimised hyperparameters. The PLP (myelin) and SMI312 (neurofilament) patches were extracted from the WM and GM boundary in the visual cortex, the Iba1 (microglia) patch was sampled from hippocampal GM, and the CD68 (activated microglia) was taken from hippocampal WM.

#### 3.2.3 Calculating the SAF map

Following segmentation, we computed SAF maps at various resolutions. First, we extracted a tissue mask by applying Otsu’s method [36] on the hematoxylin channel followed by several morphological operations. The SAF was defined as the ratio of pixels with positive DAB to those in the tissue mask within a patch. We evaluate the robustness of our pipeline by generating SAF maps at high resolution (16 × 16 μm^2^ per patch), which facilitates identification of high-frequency variations in SAF that are obscured when segmentations are pooled over larger patches. SAF maps were also generated at MRI resolution (0.5 × 0.5 mm^2^ per patch) to evaluate reproducibility at the resolution used for MRI-SAF analyses.

#### 3.2.4 Quantitative Evaluation of SAF pipeline

The SAF pipeline was evaluated with two criteria: robustness to artefacts and reproducibility of SAF estimates. These analyses used the evaluation dataset, where we assume low biological variance between adjacent slides. In cases where the protein-of-interest sparsely populates the tissue, this assumption may not always be met.

We compared the SAF maps from our pipeline with SAF maps derived manually by an expert (MPG), as described in [9]. These manually-derived SAF maps use a single dataset-specific colour matrix and segmentation threshold derived for each protein. This was achieved by manually calibrating the threshold in at least 10 randomly selected, structurally distinct regions.

To test the pipeline’s robustness to within-slide artefacts, we calculated the average column-wise SAF (Figure 6, box). These SAF profiles were created using three bandpass filters that are sensitive to different ranges of artefacts.

Within each band, we computed the relative difference in standard deviation between the manually- and automatically-derived SAF map:

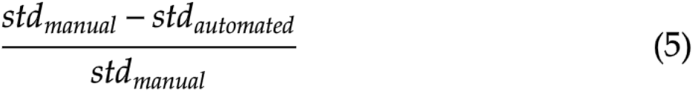

To test reproducibility of SAF estimates, we registered all within-subject IHC slides with TIRL [31] to that subject’s first slide. We calculate the absolute difference maps for each pair of co-registered slides and normalised pixelwise by the mean SAF values of both slides. The median of the difference map was calculated for each pairwise comparison of within-subject slides (Figure 7, box).

### 3.3 Co-registration of MRI and histology

After evaluation, the pipeline was applied to data from the wider multiple-region dataset (c.f. Section 2.2) which includes multiple-stain IHC and multiple-contrast MRI that was previously co-registered using TIRL (c.f. Section 2.3) [31]. The quality of co-registration was evaluated according to the alignment of tissue boundaries and WM/GM contrast. We removed data with poor quality in co-registration, where we saw substantial misalignment. In total, 3 out of 13 subjects were excluded.

### 3.4 Voxelwise correlation and regression analysis

MR parameter maps (3D) were resampled into PLP (2D) space for correlation with histology SAF. As the 2D projection of MRI voxels do not generally align with the 0.5 mm histology grid, the histology SAF were recalculated for each MR voxel, minimising interpolation effects. Pixels in the SAF maps with an intensity lower than the 5^th^ percentile were identified as non-tissue pixels and discarded. Other outliers were identified using a Huber influence function (tuning coefficient=2.5), excluding data points with weights < 0.75 [40].

For MRI-SAF analyses, we pooled voxelwise data across regions and brains for a series of linear model analyses. First, we calculated the correlation of each pair of MR (FA, MD, R2*, R1) and SAF parameters. Second, we used partial correlation to estimate the unique variance of each MR parameter that is explained by a given stain’s SAF. Subject ID was used as a covariate to account for between-subject confounds, such as post-mortem interval and fixative. We incorporated all stains and subject covariates into a single analysis to produce the final partial correlation coefficient. Finally, we performed multiple regression using all stains to derive a predictive model for each MR parameter, with subject ID as a covariate. We computed the relative importance measure [41] of each stain, i.e. the averaged relative contribution of each stain in explaining the variance of the MR parameter after the stain is added to the model.

### 3.5 Application to multiple subjects for MRI-PLP

We applied this overall workflow (Sections 3.1-3.3) to multiple subjects and brain regions in PLP slides using a total of 10 (out of 13) separate subjects (2 x healthy controls and 8 x ALS) from the same multiple-region dataset. This was after co-registration evaluation, where 3 subjects were removed due to tissue misalignment. This dataset consists of 41 PLP slides extracted from the same 3 brain regions. We correlated MRI with PLP only, as co-registration between other IHC stains and MR data was still a work-in-progress for this particular dataset. This analysis would be extremely time-consuming if conducted manually, demonstrating the value of an automated workflow. We compared the output from 10 subjects with those from the 2 subjects previously analysed (CTL 1 & ALS 1). Subject ID was modelled as a covariate.

## 4 Results

### 4.1 Quantitative Evaluation of Pipeline

To evaluate our automated pipeline, we analysed data from the evaluation dataset (Section 2.1), which includes IHC data acquired from adjacent tissue sections. Slides were processed with either the ‘default’ (Figure 2; CD68, Iba1) or ‘artefact’ (Figure 3; PLP, SMI312) pipeline. Figure 5 compares SAF maps from both our automated pipeline and a more typical ‘manual’ pipeline. Notably, both configurations of the automated pipeline reduce the impact of stitching artefacts. In the pipeline’s ‘default’ configuration, this can be attributed to the use of a slide-specific colour matrix, which results in better stain separation compared to the non-specific colour matrices used in manual analyses. The automated pipeline’s ‘artefact’ configuration also reduced staining gradient artefacts. In slides with less obvious artefacts (SMI312, Iba1), the manual and automated pipelines produce similar results.

**Figure 5:**
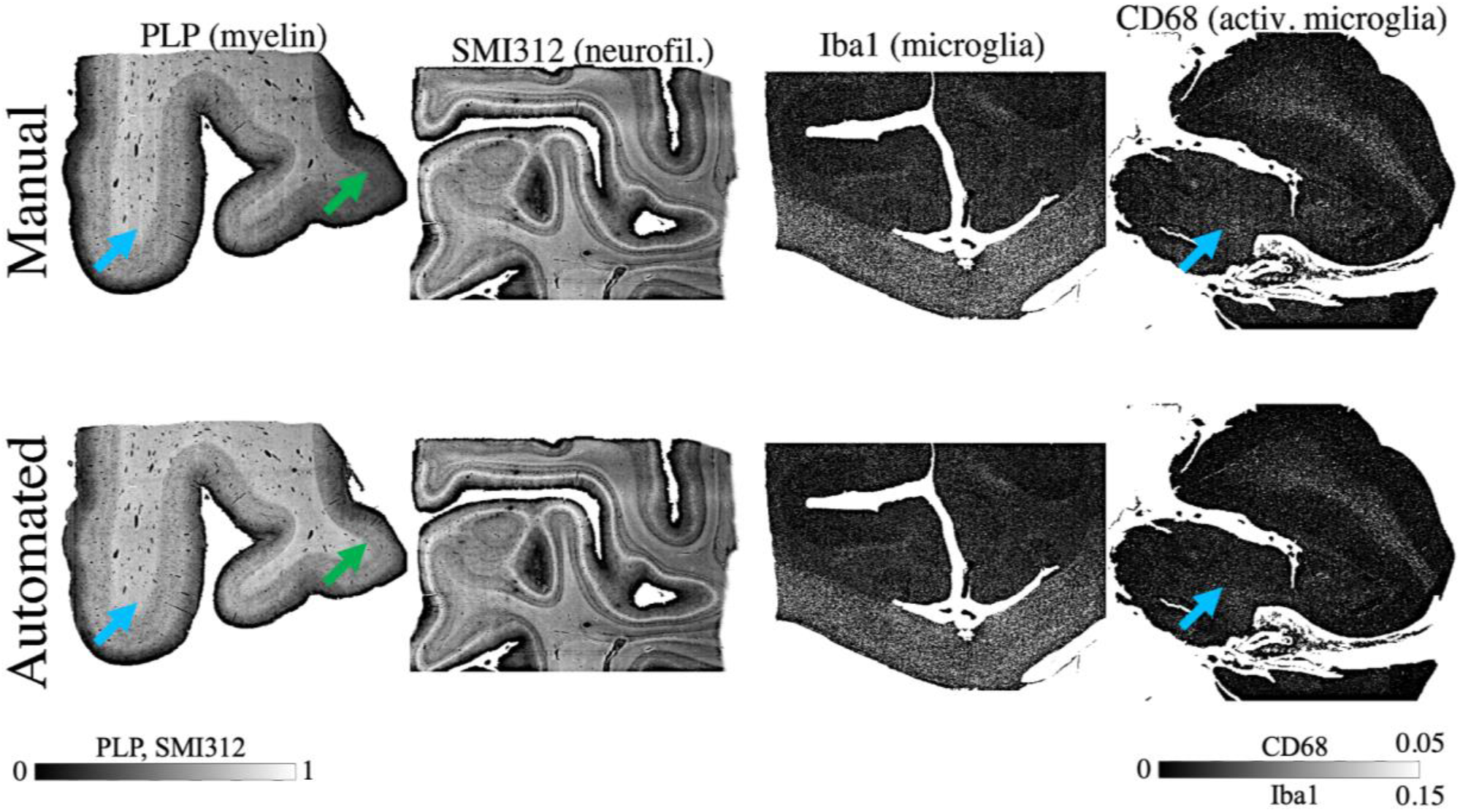
Within-slide artefacts are reduced using the automated pipeline (bottom row). These include noticeable stitching artefacts (blue arrows) and staining gradients (green arrows) originally seen in manually-derived SAF maps (top row). Examples here show PLP slides processed with the pipeline configuration that corrects for staining gradient artefacts (‘artefact’ configuration), while SMI312, Iba1 and CD68 slides were processed with the pipeline’s ‘default’ configuration.

For each slide, we quantified this reduction in artefacts via the relative percent change in standard deviation (Equation 5) of different components of the horizontal SAF profile (Figure 6 box). This horizontal SAF profile was first filtered using the lowpass and highpass filters to isolate the effects of different artefacts (staining gradients, and striping artefact with all other high frequency noise, respectively). To evaluate how much the striping artefact contributes to the overall noise, we used a bandpass filter to further isolate the frequency range of the striping artefact already separated by the highpass filter.

**Figure 6:**
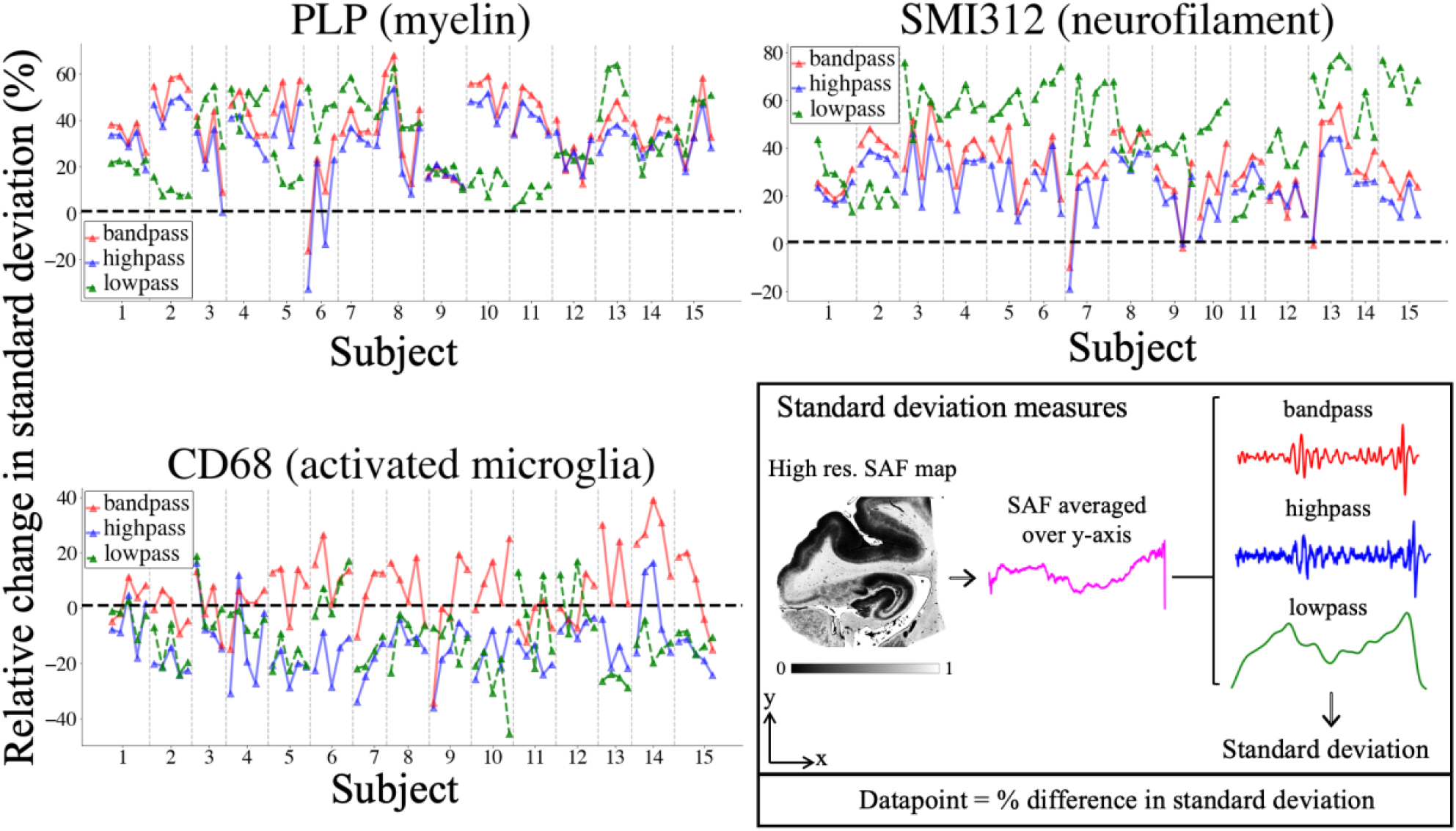
Quantitatively comparing the impact of within-slide artefacts in the manual and automated processing pipelines, as measured using the relative change in standard deviation (100 x (std_manual_ - std _automatic_) / std_manual_). Standard deviation measures (box) are derived by 1) averaging the high-resolution (16 µm/pixel) SAF map along the y-axis, 2) filtering it to produce 3 components (bandpass, highpass and lowpass), and 3) computing the components’ standard deviation. The lowpass filter isolates the staining gradient artefact, whilst the stitching artefact only is captured with the bandpass filter. The highpass filter combines the same striping artefact with all high frequency noise. Each IHC slide is represented as a single datapoint, and slides from the same subject are grouped together and connected by lines. A positive (negative) value indicates that the automatically-derived SAF map is less (more) affected by the associated artefact than the manually-derived SAF map.

In almost all PLP and SMI312 slides, the standard deviation metrics were positive, suggesting a reduction in artefacts for the automated pipeline. Notably, there was high similarity in highpass and bandpass standard deviations. This implies the striping artefact’s major contribution to overall noise in these slides, and how its impact is especially mitigated with the automated pipeline. The results for CD68 are less conclusive. The automated pipeline typically reduces the impact of the stitching artefact (the relative change within the bandpass filter is mostly positive), but performs slightly worse than the manual pipeline with respect to the highpass and lowpass filters.

Reproducibility of SAF maps (Figure 7) was also quantified by taking the median of the SAF difference map between adjacent slides from the same subject, for which true biological variability is low. Manual and automated pipelines were found to have similar reproducibility with similar variance and a median percentage change of around 5% (PLP), 9% (SMI312) and 20% (CD68). Note that fewer slides were used when evaluating CD68 (9/15 subjects; 37/73 slides) than in SMI312 (13/15 subjects; 55/72 slides) and PLP (15/15 subjects; 73/73 slides). The CD68 slides in the evaluation dataset were often observed to have inconsistent staining and/or microstructure and were excluded during manual quality control.

**Figure 7:**
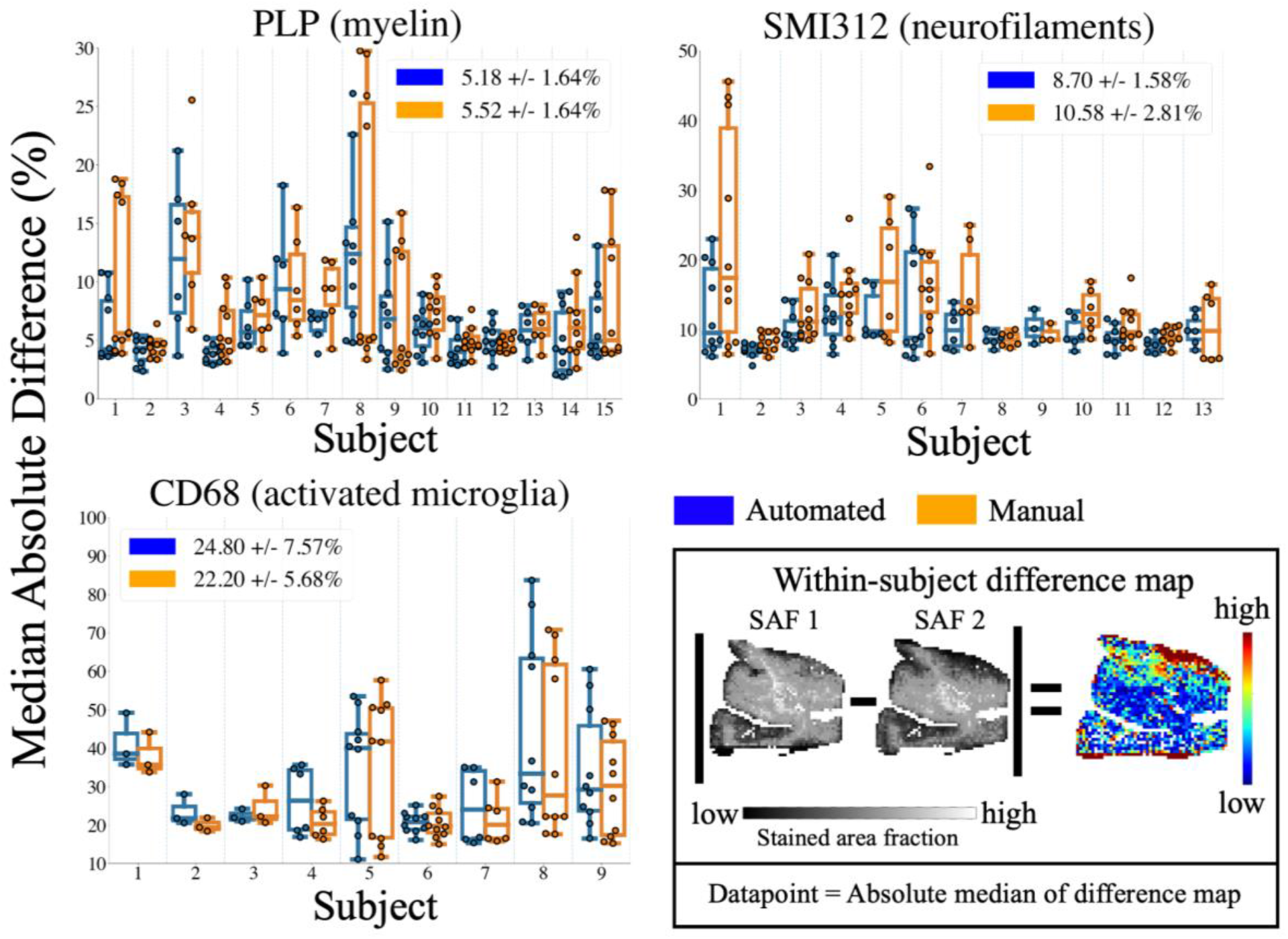
The reproducibility of SAF values across adjacent tissue sections, as measured by the median absolute difference. The median absolute difference (box) was computed by 1) taking the absolute difference of a pair of SAF maps produced from IHC slides extracted from the same region and subject, 2) normalising this difference map by the mean of both SAF maps, and 3) taking the median to represent one datapoint. We used this measure to compare reproducibility of both manually- (orange) and automatically-derived (blue) SAF maps for PLP, SMI312 and CD68. The median and median absolute variance of data points are also shown for each method (legends). Prior to reproducibility analysis, all slides were quality-checked. Slides deemed unsuitable were excluded. Note the change in scale-bar for CD68, where the difference values were generally larger when compared to PLP and SMI312.

### 4.2 Qualitative Evaluation of Co-registration

The ultimate aim for our automated pipeline is to analyse the entire multiple-region dataset. To meaningfully relate SAF map to MRI voxelwise, we require high quality MRI-microscopy co-registration. Figure 8 shows visually good alignment of both the edges of the tissue mask (green contours) and the WM/GM interface (red contour). Both contours were derived from the PLP SAF maps, given their clear WM/GM contrast. Similar results were seen in the co-registration of 10 (out of 13) other subjects (Figures S4, S5 in Appendix C).

**Figure 8:**
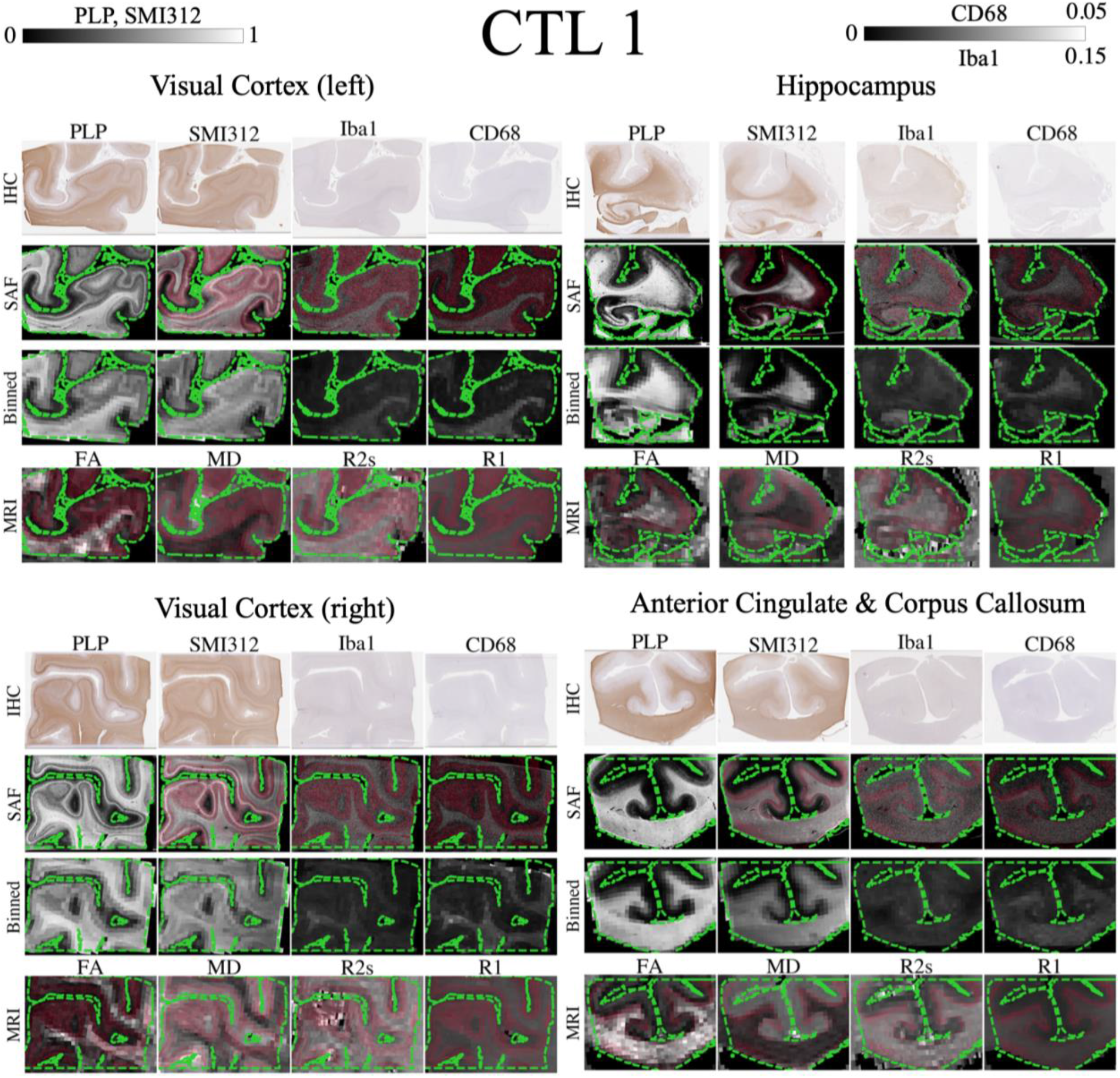
Registration evaluation for all brain regions in CTL 1. In each brain region, contours of the tissue mask (green dashed) are overlaid on the co-registered SAF maps at high resolution (first row, 1 pixel here represents SAF calculated in a 16 × 16 μm^2^ patch), SAF maps matching MRI resolution (second row) and MR parameter maps (third row). The white and grey matter interface is shown in red. The tissue boundaries are closely aligned and the high registration accuracy enables us to perform meaningful voxelwise MRI-histology correlations.

### 4.3 Correlation and regression Analysis

Voxelwise correlation and regression analysis was performed using data from two subjects in the multiple-subject study (1 x healthy control, 1 x ALS patient). Eight IHC sides corresponding to different regions (Figures 8, S4) were mapped onto MRI data. For each MR parameter, up to 3% of voxels were classified as outliers and removed, resulting in an average sample size of 6506 voxels (FA=5846, MD=5852, R2*=8956, R1=5373). Of these non-outlier voxels, an average of 2246 voxels were classified as WM (FA=2039, MD=2029, R2*=3041, R1=1877).

#### 4.3.1 Univariate linear regression of MRI and SAF

Univariate linear regression was used to relate each MRI-SAF pair (Figure 9). Voxels from both WM and GM were pooled across brain regions and subjects. While correlations with Iba1 appear low (|r|=0.035-0.28), CD68 correlated well with FA (r=0.56) and MD (r=-0.39). The scatter plots also suggest nonlinear trends for FA with PLP and SMI312. Other tissue features that are known to alter FA, most notably fibre dispersion, are not accounted for in this study. These features may correlate to myelin (PLP) or neurofilament (SMI312) fraction, and thus relate to FA. A similar scatterplot using voxels from WM only is shown in Figure S3 (Appendix C).

**Figure 9:**
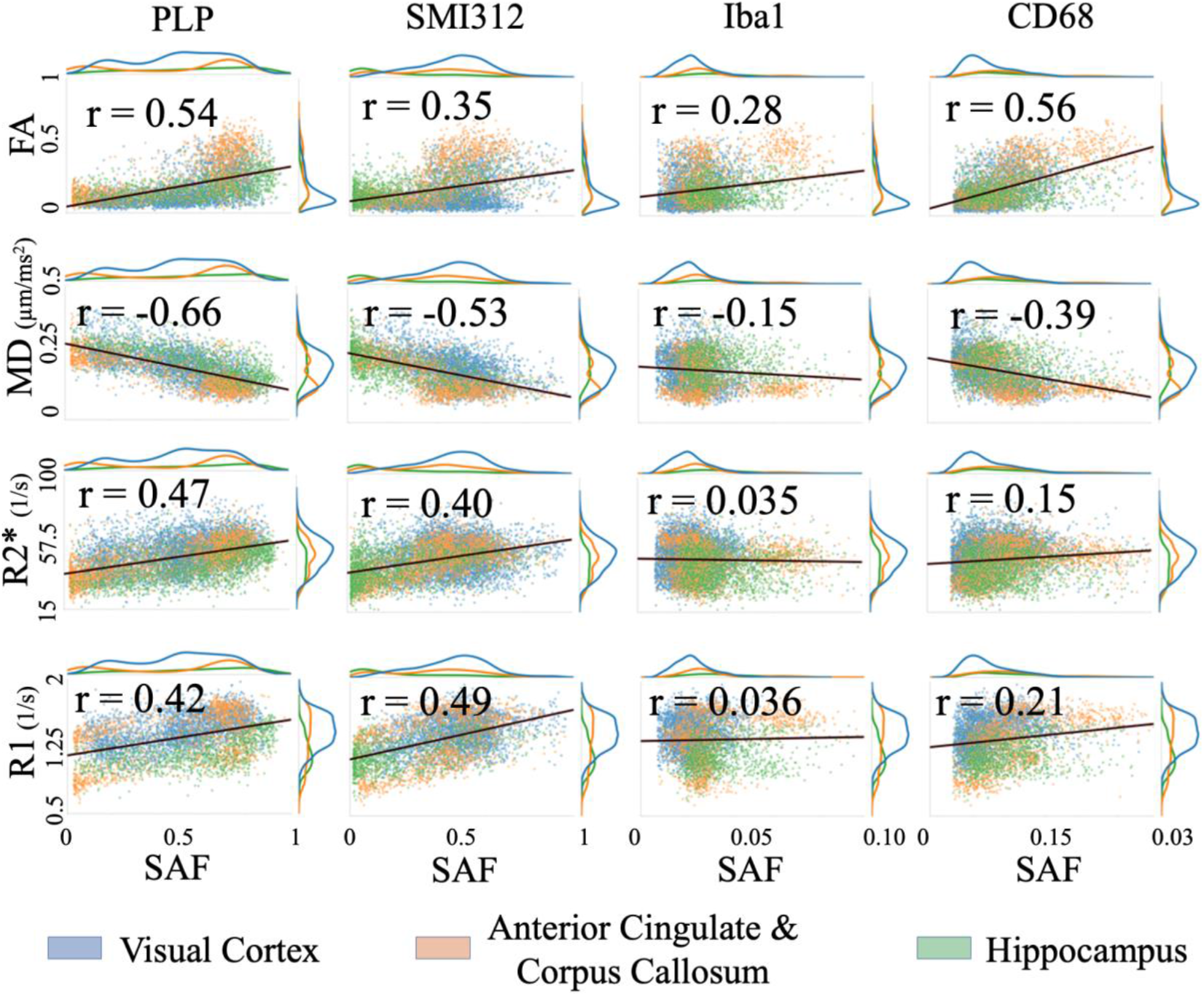
Correlating MR parameters (DTI FA, MD, R2* and R1) with IHC SAF. The line of best fit (black line) and corresponding Pearson correlation coefficients, r, are overlaid. The visual cortex (blue), anterior cingulate (orange) and hippocampus (green) provide good dynamic range for the MR parameters and SAFs.

#### 4.3.2 Partial correlations of MRI with IHC

Partial correlations were used to investigate the unique variance of each MR parameter that is explained by an IHC stain. For a given stain, a series of correlations were calculated after regressing out one other individual stain or all other stains as covariates. We show correlation coefficients when including all voxels (WM,GM) (Figure 10), or WM only (Figure 11).

**Figure 10:**
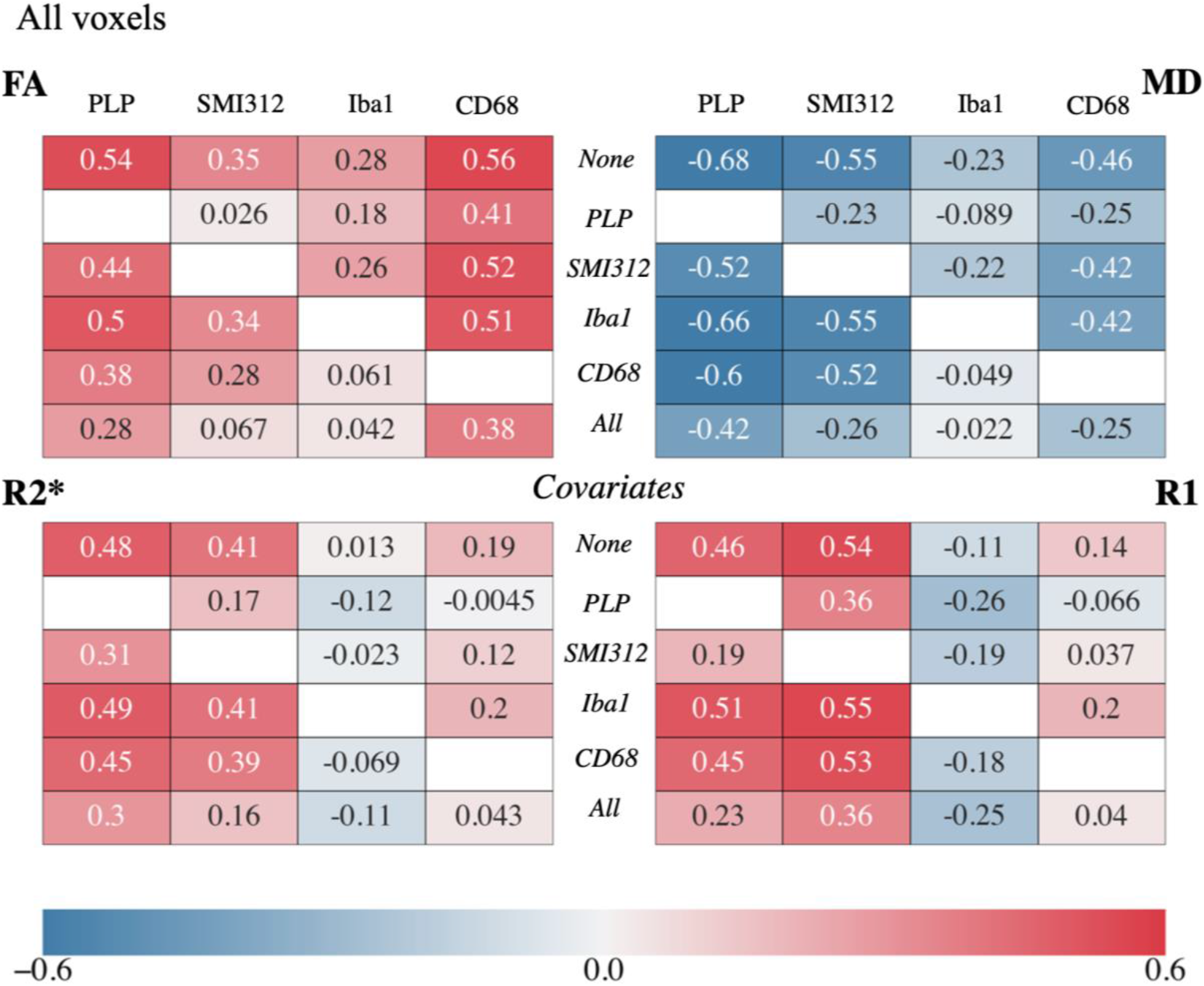
Partial correlation analysis between MR parameters (FA, MD, R2* and R1) and IHC stains for both white and gray matter voxels (Fig. 12 shows results for white matter voxels only). In each quadrant, the top row (“None”) gives the correlation coefficients when accounting for subject ID only. The middle rows give the partial correlation coefficient controlling for one of the other stains and subject ID (italicised labels in the second to fifth rows). The bottom row (“All”) gives the partial correlation coefficient controlling for all stains and subject ID. *PLP (myelin); SMI312 (neurofilaments); Iba1 (microglia); CD68 (activated microglia)*.

**Figure 11:**
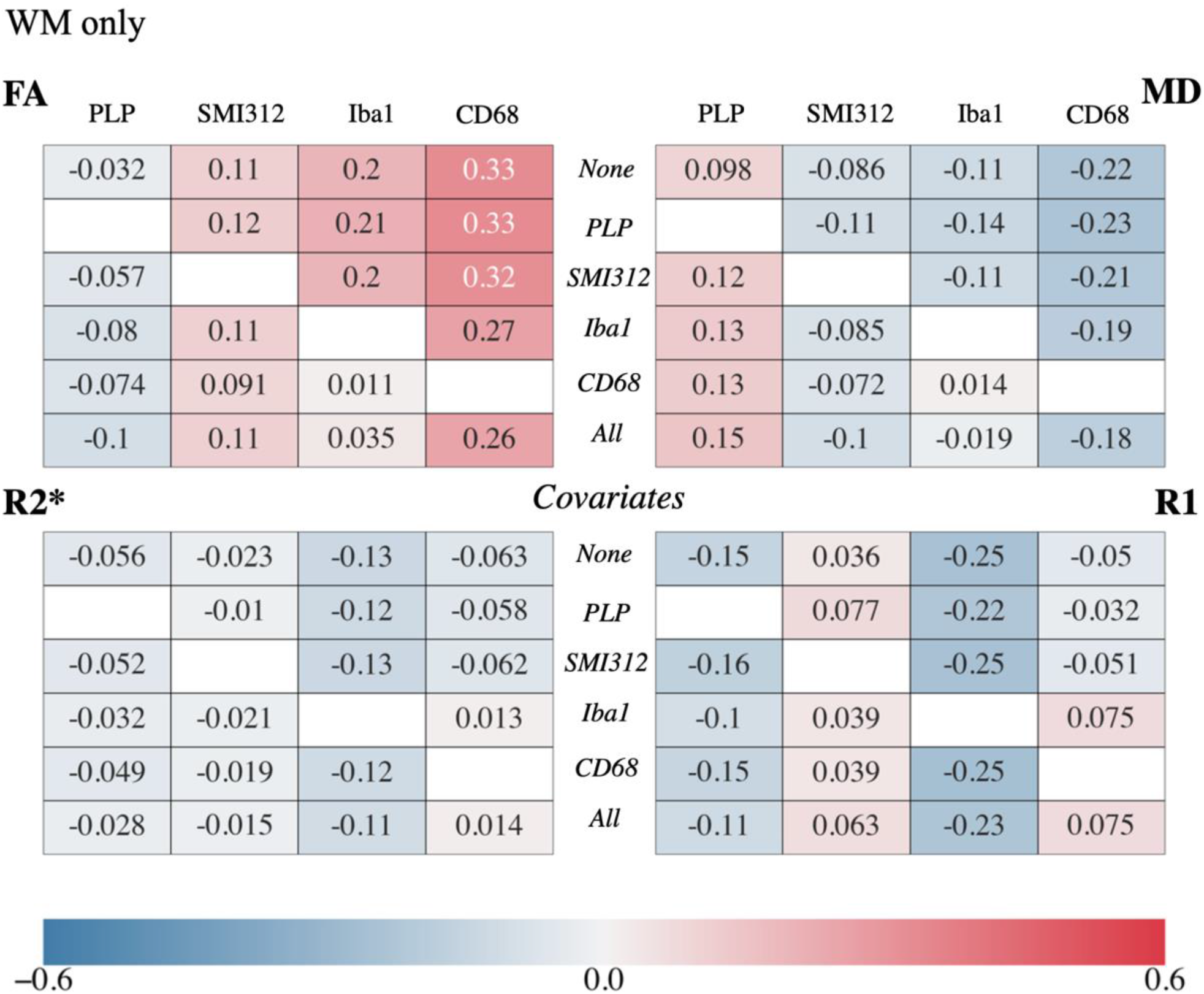
Partial correlation analysis between MR parameters and IHC stains for voxels from the WM only (Figure 10 shows results for all brain voxels). The interpretation is similar to Figure 10. Note that correlations between FA and MD with CD68 (activated microglia) remain relatively high, even after accounting for all other stains. High correlations also remain between R1 and R2* with Iba1 (microglia).

##### WM and GM

When accounting for all other stains (Figure 10, bottom rows), FA is best explained by CD68 (r=0.38), MD by PLP (r=-0.42), R2* by PLP (r=0.30) and R1 by SMI312 (r=0.36). We observed two main results: the shared explained variance between PLP (myelin) and SMI312 (neurofilaments), and between Iba1 (all microglia) and CD68 (activated microglia).

First, the correlation coefficient between FA and SMI312 is greatly reduced after accounting for PLP (r=0.35 to 0.026), whilst the correlation coefficient between FA and PLP is minimally reduced after accounting for SMI312 (r=0.54 to 0.44). Similar effects are observed between MD and PLP/SMI312, and between R2* and PLP/SMI312. Together, this suggests that there is unique variance in FA, MD and R2* explained by myelin in particular, but that associations with neurofilaments may primarily reflect a spatial covariance with myelin rather than a direct relationship. The opposite effect is seen in R1: the correlation coefficient between R1 and PLP decreases significantly when covarying for PLP (r=0.46 to 0.19), but the correlation coefficient between R1 and SMI312 is marginally reduced when accounting for PLP (r=0.54 to 0.36). This implies that R1 is sensitive to neurites in general, rather than myelin itself. Second, we saw that the high correlation of FA with CD68 only marginally decreases after accounting for Iba1 (r=0.56 to 0.51). Conversely, the apparent correlation with Iba1 is minimal after accounting for CD68 (r=0.28 to 0.061). We note similar trends for correlations of MD with CD68/Iba1.

##### WM only

Figure 11 shows the same analysis for voxels found in the WM only. When accounting for all other stains, FA was best explained by CD68 (r=0.26), MD by CD68 (r=-0.18), R2* by Iba1 (r=-0.11) and R1 by Iba1 (r=-0.23). When compared to our previous results for WM and GM voxels, the correlation coefficient between the four MR parameters and PLP/SMI312 are greatly reduced and/or close to zero. This suggests that the correlations of MR parameters with PLP/SMI312 are primarily driven by WM/GM contrast. This behaviour is not seen for CD68 and Iba1. When considering WM voxels only, CD68 explains the most unique variance with FA. Notably, this correlation remains relatively unchanged when accounting for other stains (r= 0.33 to 0.26). Similar behaviour is observed when relating CD68 with MD, and Iba1 with R1/R2*.

#### 4.3.3 Multiple linear regression of MRI with IHC

Multiple linear regression was performed to derive a predictive model for each MR parameter that is driven by multiple stains. Each MR parameter was modelled as a linear combination of all stains (explanatory variables) and the subject ID (confounding variable). We show the regression and the fitted correlation coefficients (i.e. square root of the coefficient of determination) when including voxels from both WM and GM (Table 1A), or WM only (Table 2A). Using multiple stains better explains the variance in MR parameters than individual stains. We also show the relative importance of each predictor in influencing MR parameters (Tables 1B, 2B). Notably, the “subject” predictor had the highest relative importance in predicting R1 in all voxels, and MD/R2*/R1 in WM voxels only. This implies that between-subject confounds, due to the known effects of sample handling factors like post-mortem interval, substantially influence MR parameters.

**Table 1:** Multiple linear regression predicting MR parameters using multiple IHC stains. A “subject” variable was also included to account for confounds such as post-mortem interval and age effects. As the predictors are unitless, all offsets and regression slopes are given in units of MR parameters. A: The regression coefficients and correlation coefficients r_fit_. B: The relative importance of each stain describes the amount of variance it can explain in an MR parameter, averaged over all permutations of multiple regressions that include the specific stain. Values are normalised across stains to get a unit percentage.

| | Offset | PLP | SMI312 | Iba1 | CD68 | Subject | $r_{\text{fit}}$ |
| --- | --- | --- | --- | --- | --- | --- | --- |
| FA | -0.00142* | 0.173 | 0.0429 | 0.252 | 10.5 | -0.0112 | 0.645 |
| MD<br>[$\mu\text{m}^2/\text{ms}$] | 0.286 | -0.112 | -0.070 | -0.0629* | -2.55 | 0.0366 | 0.745 |
| R2*<br>[1/s] | 39.2 | 16.5 | 9.04 | -61.2 | 80.7 | -3.73 | 0.536 |
| R1<br>[1/s] | 1.04 | 0.220 | 0.382 | -2.48 | 1.34 | 0.234 | 0.710 |

\*not significant ( $p > 0.05$ )

**Table 1:** Multiple linear regression predicting MR parameters using multiple IHC stains.
|  | PLP | SMI312 | Iba1 | CD68 | Subject |
| --- | --- | --- | --- | --- | --- |
| FA | 36.0 | 11.9 | 7.04 | 44.5 | 0.55 |
| MD | 45.0 | 25.7 | 1.69 | 14.5 | 13.1 |
| R2* | 51.3 | 30.1 | 2.90 | 3.86 | 11.8 |
| R1 | 18.7 | 30.6 | 3.15 | 2.93 | 44.6 |

**Table 2:** Multiple linear regression now computed from voxels located in the tissue’s WM. These tables are interpreted similar to Table 1.

| | Offset | PLP | SMI312 | Iba1 | CD68 | Subject | $r_{\text{fit}}$ |
| --- | --- | --- | --- | --- | --- | --- | --- |
| FA | 0.255 | -0.173 | 0.144 | 0.410* | 8.81 | -0.0247 | 0.353 |
| MD<br>[$\mu\text{m}^2/\text{ms}$] | 0.118 | 0.0610 | -0.0323 | -0.0531* | -1.40 | 0.0310 | 0.429 |
| R2*<br>[1/s] | 61.2 | -2.22* | -0.889* | -55.2 | 19.6* | -5.54 | 0.455 |
| R1<br>[1/s] | 1.52 | -0.217 | 0.0878 | -2.71 | 1.89 | 0.268 | 0.637 |

\*not significant ( $p>0.05$ )

|  | PLP | SMI312 | Iba1 | CD68 | Subject |
| --- | --- | --- | --- | --- | --- |
| FA | 4.32 | 8.75 | 15.1 | 69.2 | 2.66 |
| MD | 6.43 | 2.40 | 4.99 | 16.2 | 70.0 |
| R2* | 0.47 | 1.97 | 22.6 | 3.62 | 71.3 |
| R1 | 4.54 | 2.39 | 5.88 | 0.443 | 86.8 |

### 4.4 Application to multiple subjects for MRI-PLP

The primary benefit of the pipeline is that it can be rapidly applied across many subjects, which we demonstrate by applying to ten other subjects (2 x healthy control, 8 x ALS patients, 41 slides) from the larger dataset to perform a similar univariate MRI-PLP linear regression analysis. Note that these ten subjects are separate from the two subjects used for multiple linear regression analysis. Following outlier removal, we had an average of 31,047 voxels/MR parameter (FA=28,849, MD=28,598, R2*=44,134, R1=22,608). In WM, we had an average of 10,724 voxels/MR parameter (FA=10,040, MD=9954, R2*=14,832, R1=8072). Similar trends and regression coefficients were seen for all parameters for the larger group (n=10, Figure 12 right) as were found in the original analysis (n=2, Figure 12 left), with the estimated effect (**β**) being highly consistent for FA, MD and R2*, but less consistent for R1.

**Figure 12:**
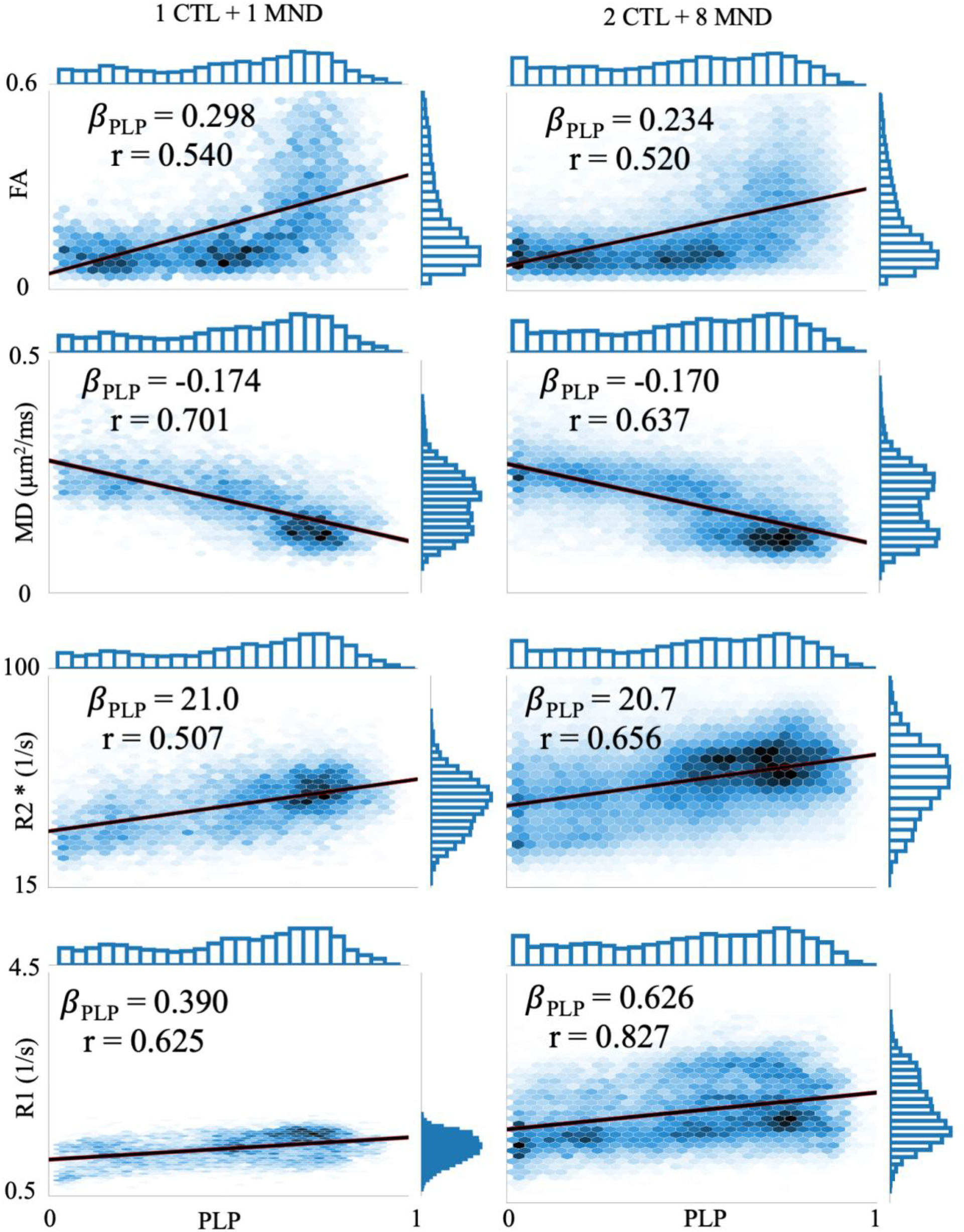
Density plots showing the relationship of different MR parameters (rows) with PLP (myelin). We compare the MRI-PLP regression coefficient (***β***_***PLP***_) derived from analysis across many subjects (CTL 2-3 & ALS 2-9; right column) with analysis across 2 subjects (CTL 1 & ALS 1; left column). The effect size, ***β***_***PLP***_, is highly consistent between the two datasets for FA, MD and R2*, but less so for R1.

## 5 Discussion

To facilitate high-throughput MRI-histology analyses, we introduce an automated pipeline to extract a quantitative metric, stain area fraction (SAF), from immunohistochemical stains. Using high-quality co-registration, we performed whole-slide voxelwise MRI-SAF comparisons. The pipeline was applied to post-mortem human brain data from multiple subjects, relating IHC slides stained for myelin (PLP), neurofilaments (SMI312), microglia (Iba1) and activated microglia (CD68) to MR parameters (FA, MD, R2*, R1).

Compared to previous literature, our approach has three advantages. First, while most MRI-SAF studies use manually-derived SAF, our pipeline deploys data-driven algorithms [34, 36]. This enables the pipeline to be applied rapidly to larger datasets without expert intervention. This is demonstrated in how we applied the pipeline to 41 myelin-stained slides, with each slide taking approximately 2 hours. Second, our pipeline modifies these automated algorithms to handle key non-biological sources of variation, which otherwise lead to artefacts in the SAF maps. This is an improvement over previous automated pipelines [42, 43, 44]. Third, the pipeline is generalisable to multiple IHC stains, allowing for analysis spanning multiple microstructural sources. Combined with recent advances in MRI-microscopy co-registration [31], the pipeline may enable standardised analyses for voxelwise MRI-SAF comparisons [14].

After running the pipeline, we perform voxelwise MRI-SAF analysis. This inspires more confidence in the derived MRI-SAF correlations compared to ROI-based analyses, which may be biased by the choice of ROI and/or dilute localised effects-of-interest. Further, most studies correlate MRI with only a single, or a few histological markers [10, 18, 23, 42, 45]. Here, we relate multimodal MRI with multiple IHC stains. This facilitates more complex analyses, such as partial correlation, which elucidate how each microstructure might contribute to MRI signal changes while accounting for other colocalised microstructure.

### 5.1 SAF Pipeline

We evaluated the SAF pipeline to determine 1) its robustness to IHC artefacts, alongside 2) its ability to produce reproducible SAF value. Here, we analysed a separate evaluation dataset which included 4-5 adjacent slides (per stain) sampled from each subject. We used this dataset to separate sources of biological variability, which is assumed to be minimised in adjacent slides, from SAF variations associated with tissue processing or IHC analysis.

Our results show the pipeline’s robustness to common IHC artefacts, while maintaining similar reproducibility to manually-derived SAF values (Figures 5-7). These artefacts were particularly prominent in densely-stained slides such as PLP, which almost always had observable within-slide artefacts. Our pipeline reduced these artefacts, as seen in the positive standard deviation metrics for most slides.

Conversely, CD68 appears less reproducible for both manual and automated processing. CD68 stains sparse, small activated microglia (soma diameter∼10 µm [46]), for which we expect higher biological variability between adjacent slides This may make the SAF intrinsically less reproducible. Considering within-slide artefacts, staining gradients were not visibly present in CD68 slides, although some striping was visible on the manually-derived SAF maps. The automated pipeline reduced this striping, as reflected in the improved bandpass standard deviation metric. While many CD68 slides were excluded from the reproducibility dataset due to IHC protocol non-idealities that affect batches of slides no CD68 slides were excluded during QC of the separately acquired multiple-region dataset.

### 5.2 MRI-SAF analysis

The multiple-region dataset includes MRI and IHC data that has been carefully co-registered using TIRL, facilitating meaningful voxelwise analysis. Our correlation results agree with previous studies for myelin [14, 16, 18, 23] and neurofilaments [42]. Conversely, few studies report MRI-microglia correlations in the human brain. Our results for microglia correlate positively with FA and negatively with MD, similar to that reported in the human spinal cord [42].

#### 5.2.1 Myelin and neurofilaments

Our partial correlations for PLP (myelin) and SMI312 (neurofilaments) demonstrate pitfalls in performing pairwise MRI-histology correlations without controlling for other tissue features. This is illustrated in how there is unique variance in FA/MD/R2* explained by myelin co-localized with neurofilament, rather than neurofilament itself. The opposite is shown from correlations with R1. This is noteworthy, given the studies correlating myelin with R1 [21, 47-49] while not controlling for neurofilaments. In WM only, PLP/SMI312 have reduced associations with all MR parameters (Figure 11), suggesting that these associations were being primarily driven by WM/GM contrast. Taken together, our results may help explain the large range of MRI-myelin correlation coefficients reported in two reviews (Figures 4, 5 in [14]; Figure 4 in [16]).

#### 5.2.2 Microglia

Our results suggest a relationship between CD68 (activated microglia) and FA/MD, even after accounting for Iba1 (all microglia). This relationship is present when considering all voxels (WM, GM), and WM only. Activated microglia are a generic biomarker of inflammation [50, 51], implying a sensitivity of DTI measures to neuroinflammation. Intriguingly, in-vivo studies report an opposite trend, where FA decreased with alternate markers of neuroinflammation in Parkinson’s disease [52] and obesity [53]. However, the use of different markers makes comparison with our results difficult. Future experiments may investigate how activated microglia affects diffusion-weighted MRI by using more sophisticated biophysical models [54, 55].

The correlations of R1/R2* with Iba1 remained high after accounting for CD68, suggesting that net microglial load relates to relaxometry measures. Surprisingly, microglia SAF correlates negatively with R1/R2*, whereas iron is known to colocalise with activated microglia [56, 57] and correlate positively with R1/R2* [22, 23, 47, 59]. Future studies will need to incorporate iron-stained IHC to elucidate the contributions of iron to relaxometry measures.

### 5.3 Limitations

There are limitations to this study. First, we use SAF to quantify MRI-histology relationships, which may not scale linearly with protein density. While our results show key trends between MR parameters and proteins, these MRI-SAF slopes may not be directly comparable to other studies. Second, we do not incorporate other histological metrics that may relate to MRI. These include the fibres’ orientation dispersion and microglial cell morphology, which may explain variance in MRI. Finally, our pipeline does not address smaller-scale within-slide artefacts, such as artifactual staining of vasculature and folding artefacts at tissue edges [26]. These might bias estimated SAF. Since our analysis occurs at the slide-level, manually excluding all smaller-scale artefacts would be very time-intensive. Given the high number of analysis data points, we expect the effect size of these artefacts to be small.

## 6 Conclusion

We introduce an automated pipeline that extracts SAF maps from IHC slides. By design, the pipeline is generalisable to multiple IHC stains and robust to artefacts from tissue staining and/or slide digitisation. The pipeline was applied to co-registered MRI and IHC data from post-mortem human brains, where we perform multiple linear analyses to investigate voxelwise relationships between MRI- and IHC-derived parameters (FA, MD, R1 and R2* against SAF for myelin, neurofilaments and microglia). Our results emphasise the need to simultaneously analyse multiple stains when validating MRI, as to avoid misleading inference due to the spatial covariance of multiple microstructural features. Interestingly, we found several diffusion-weighted metrics’ sensitivity to activated microglia – a biomarker of neuroinflammation. Together, our results demonstrate the pipeline as a valuable tool for IHC analyses and future investigations relating MRI to disease pathology.

## Availability of code and data

The code for this work is publicly available at https://git.fmrib.ox.ac.uk/spet4877/ihcpy. The data will soon be available on https://open.win.ox.ac.uk/DigitalBrainBank/.

## Authors’ Contributions

**DZLK** conceived the study design, developed the histology processing pipeline, implemented the MRI-histology workflow, conducted data analyses, and drafted the manuscript. **SJ** conceived the study design, contributed to the design of the MRI-histology workflow and data analyses. **INH** developed the TIRL registration platform, performed the MRI-histology co-registration, designed the acquisition of the reproducibility dataset, and advised on data analyses. **JM** contributed to the histology processing pipeline. **BCT** analysed the diffusion MRI and T1 data. **SF** developed the MRI protocols and acquired the MRI data. **CW** analysed the R2* data. **CS** contributed to the acquisition of the multiple-region dataset. **AS** performed the histological processing, curated histological data, and provided expert feedback on histology analysis. **OA** designed the brain sampling strategy and provided expert feedback on histological analysis. **MPG** optimised histological protocols, designed the brain sampling strategy, performed histology analysis, evaluated histological segmentations, and provided expert feedback on histological analysis. **KLM** conceived the study design, designed the post-mortem MRI protocol, contributed to the MRI-histology workflow, contributed to all data analysis, and helped draft the manuscript. **AFDH** conceived the study design, contributed to the MRI-histology workflow, contributed to all data analysis, and helped draft the manuscript.

## Declaration of Competing Interests

None

## Acknowledgements

AFDH and KLM contributed equally to this work. Data acquisition was funded by the grant MR/K02213X/1 from the Medical Research Council (MRC). DZLK is supported by the Hrothgar Singaporean Clarendon Scholarship and the Nuffield Department of Clinical Neurosciences studentship. AFDH and INH are supported by the EPSRC and MRC grants (grants EP/L016052/1 and MR/L009013/1). KLM, AFDH, INH, BCT, CW, OA and SJ are supported by the Wellcome Trust (grants WT202788/Z/16/A, WT215573/Z/19/Z and WT221933/Z/20/Z). The Wellcome Centre for Integrative Neuroimaging is supported by core funding from the Wellcome Trust (203139/Z/16/Z). The authors are grateful to the donors and benefactors of the Oxford Brain Bank that kindly provided all human tissues for this study.

## Supplementary Materials

## Appendix A. Immunohistochemistry protocol

After post-mortem MRI, tissue samples of key regions were extracted as described in [9]. Tissue samples were processed and embedded in paraffin. 6-µm thick paraffin sections were cut and mounted on 75 × 26 mm slides for immunohistochemistry. Sections were then deparaffinised in xylene and rehydrated in graded ethanol and water. Endogenous peroxidase activity was blocked by 3% H_2_O_2_ (in phosphate buffered saline) for 30 minutes before carrying out heat-induced antigen retrieval in citrate buffer (pH=6). Sections were incubated with primary antibodies against SMI312 (1:2000, Biolegend 837901), PLP (1:1000, Bio-Rad MCA839G), Iba1 (1:1000, WAKO 019-19741) and CD68 (1:50, DAKO M0876) for 1 hour. These proteins were visualised with DAKO EnVision Detection Systems. Antibody horseradish peroxidase rabbit/mouse serum was applied for 1 hour followed by 3,3’-diaminobenzidine (DAB) chromogen. Sections were counterstained with haematoxylin, dehydrated with graded ethanol, cleared in xylene and mounted with DPX mounting medium.

## Appendix B. Colour Channel Separation

The stain separation technique used to compute a stain density uses Beer-Lambert’s law. In Beer-Lambert’s law, absorbance (A_λ_) is used to relate the attenuation of monochromatic incident light (I_inc,λ_) passing through the IHC slide to the detected light (I_det,λ_):

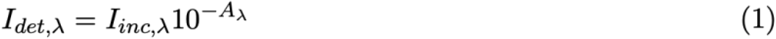

where A_λ_:

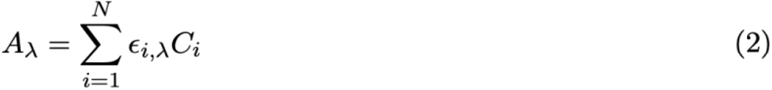

Here, *N* represents the number of stains, ε_i,λ_ denotes the attenuation coefficient of i^th^ stain for light at wavelength λ, and C_i_ represents the stain density of i^th^ stain. For λ corresponding to red, green and blue colour channels, the stain density (**C**_**stain**_) of DAB, hematoxylin, and the leftover staining (residual) for a pixel with I_det,λ_ on the slide can be derived as earlier presented:

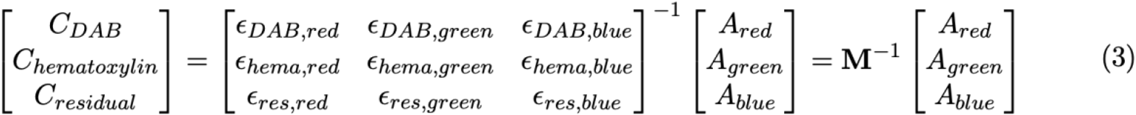

With the derivation of a colour matrix **M**, estimation of stain density, **C**_**stain**_, is a least-squares error minimization problem solved by multiplying absorbance, **A**_**R**,**G**,**B**_, values with an inverse of M. However, the rows of M (i.e. the stain-specific colour vectors) are not constrained to be orthogonal. This means that the representation of some valid colour vectors requires estimation of **C**_**stain**_ that are negative. While this is mathematically valid, it leads to a problematic physical interpretation of C_stain_ in terms of actual stain concentrations (i.e. that a stain actually emits, rather than absorbs, light).

We address this by using a non-negative approach designed to emulate the non-negative least squares (NNLS) algorithm. We call this the pseudo-NNLS (pNNLS). The pNNLS is used over NNLS as it computationally much faster than the NNLS. The pNNLS first estimates a stain’s density C_stain_ via a standard matrix inversion. In the non-physical case where a stain’s C_stain_ is negative, the stain’s density value is set to 0. The other stain’s positive density value is then re-computed by projecting the **AR**,**G**,**B** to the specific stain’s colour vector in isolation. We conducted simulations (not shown) to confirm that stain densities derived from both pNNLS and NNLS are effectively identical. Note that since all A_R,G,B_ values are strictly positive, the case of having negative densities for both stains simultaneously is not possible.

## Appendix C. Figures

**Figure S1:**
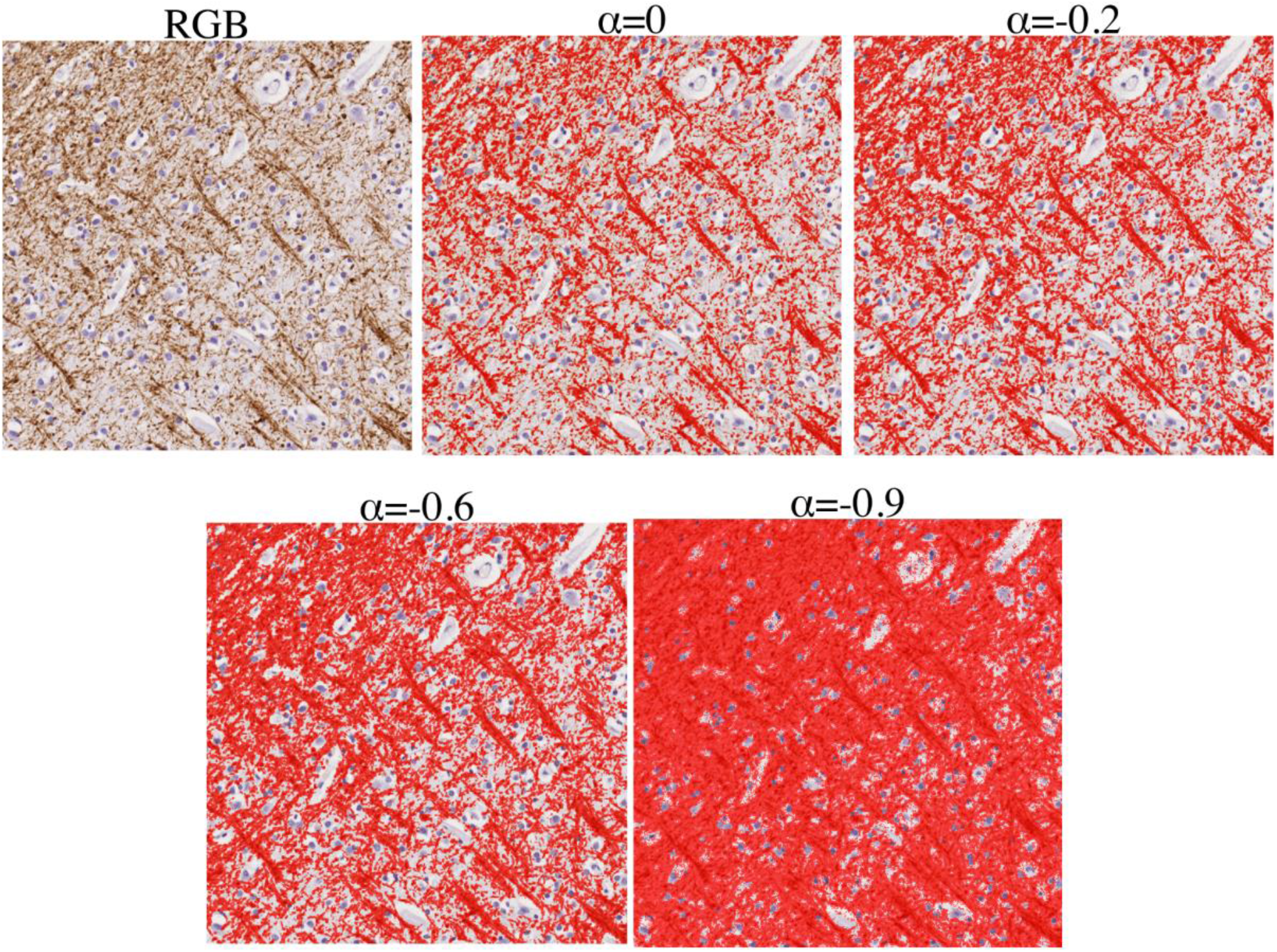
Optimising hyperparameter **α** to produce an accurate segmentation of the target protein. We show this in an example myelin-stained (PLP) patch. The hyperparameter α affects the segmentation of the protein-of-interest. Here, varying **α** from 0 to −0.9 slowly increases the number of pixels we identify as protein-of-interest. Based on expert input, we identified **α**=-0.6 as optimal for accurately segmenting myelin. Once **α** is set for a single stain, we apply it to all slides of that stain.

**Figure S2:**
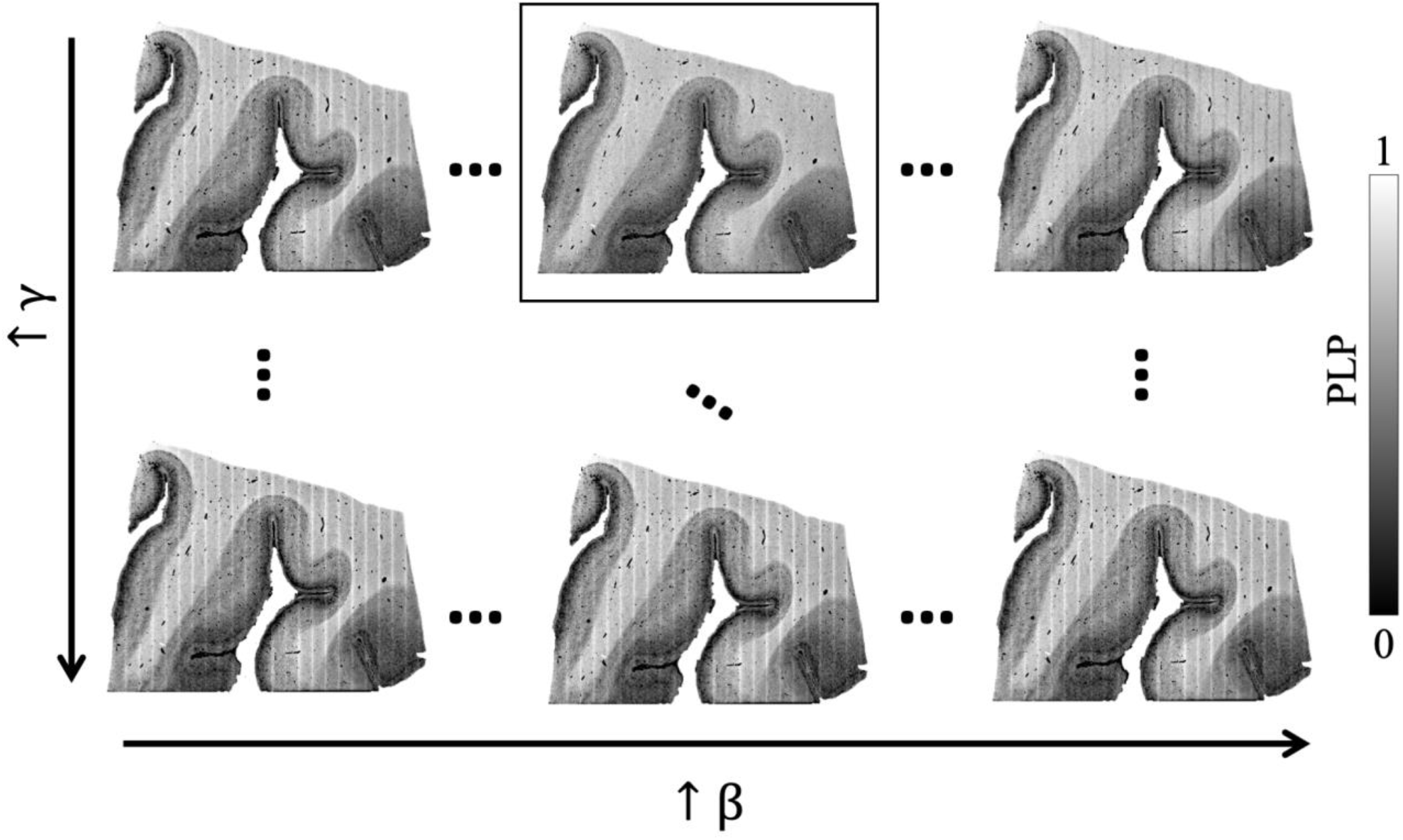
Optimising hyperparameters **β, γ** to produce a resultant stain area fraction map. These hyperparameters directly address the striping artefacts, and are automatically determined for each slide. In this example myelin-stained (PLP) slide, we see substantial striping in the SAF map (top left; **β**=0; **γ**=0), An increase in **β** is needed to produce segmentation threshold that compensate for changes due to striping present (black square; **β**=1.5, **γ**=1), though a **β** which is too high may also result in an overcompensation (top right; **β**=4, **γ**=1). Finally, hyperparameter **γ** is a smoothing kernel applied to the segmentation thresholds, to prevent the thresholding from varying abruptly from column-to-column. If sharper striping is observed, a smaller **γ** is used. Conversely, over-smoothing these segmentation thresholds (high **γ**) may also effectively negate adjustments to the thresholds made by **β** (last row).

**Figure S3:**
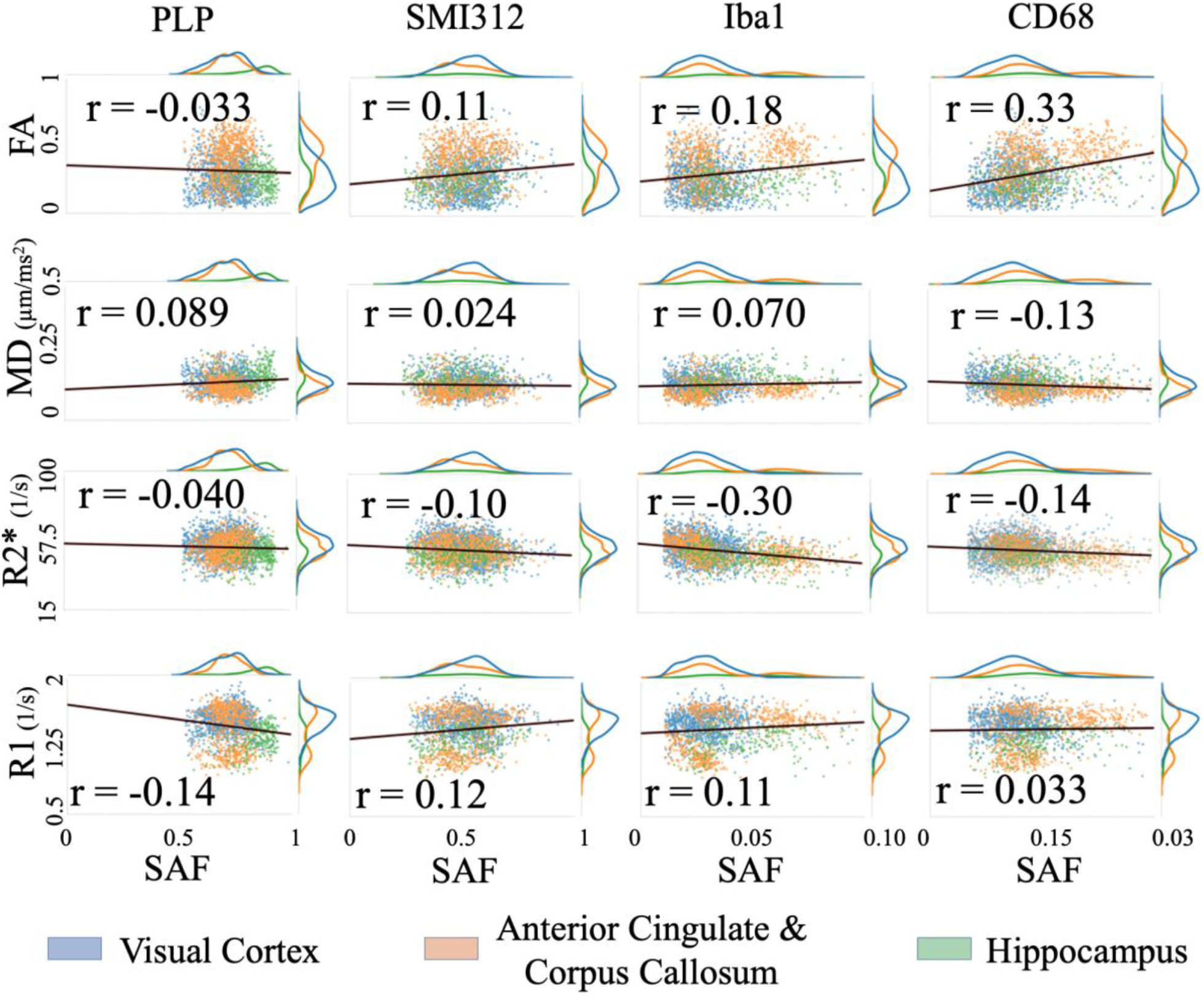
Correlating MR parameters (DTI FA, MD, R2* and R1) with IHC SAF in white matter only. The line of best fit (black line) and corresponding Pearson correlation coefficients, r, are overlaid. Again, the visual cortex (blue), anterior cingulate (orange) and hippocampus (green) provide good dynamic range for the MR parameters and SAFs.

**Figure S4:**
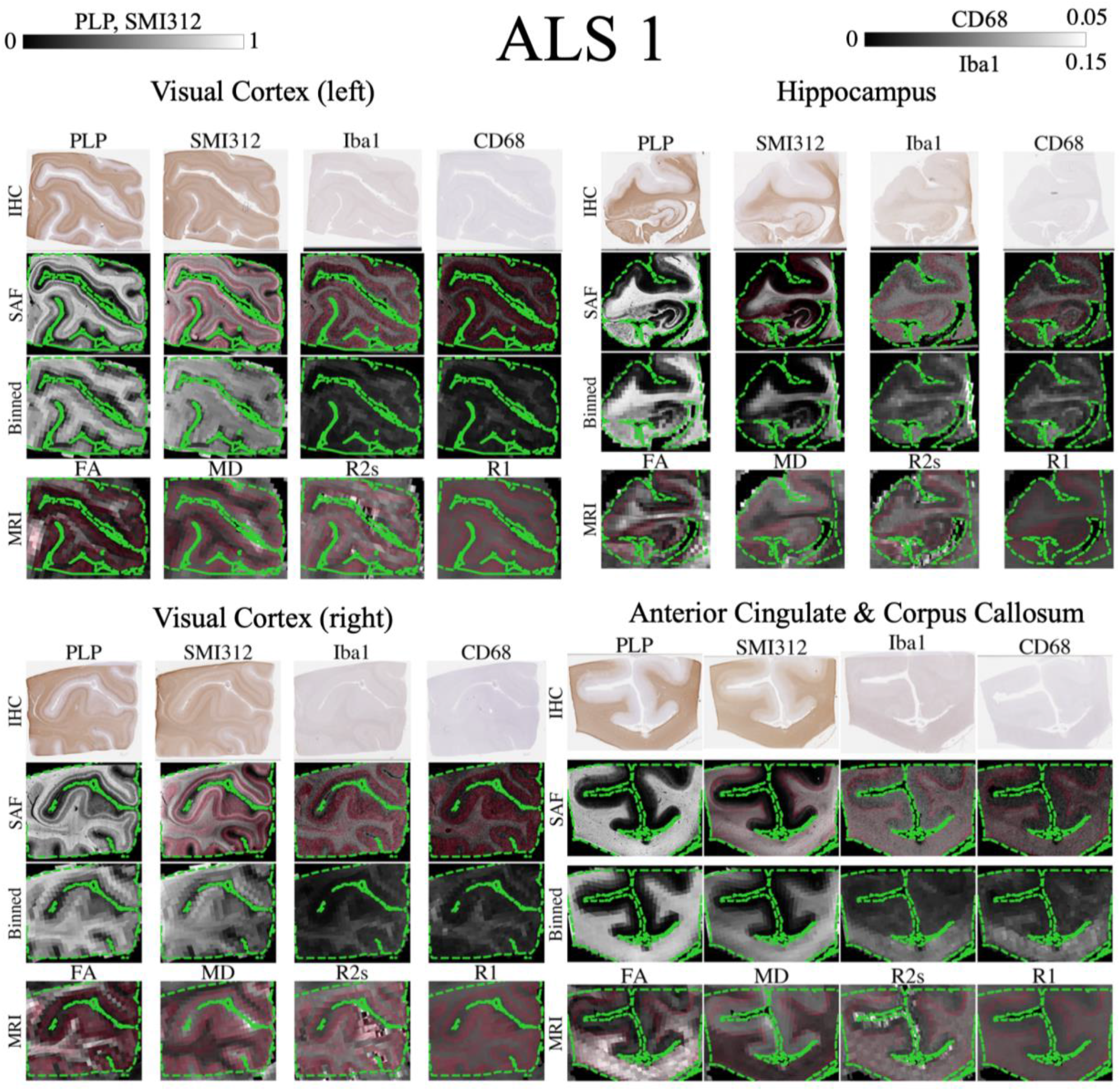
Registration evaluation for all brain regions in ALS 1. Results are displayed as previously described in Figure 8. Again, we see close alignment of tissue borders and contours representing WM/GM contrast.

**Figure S5:**
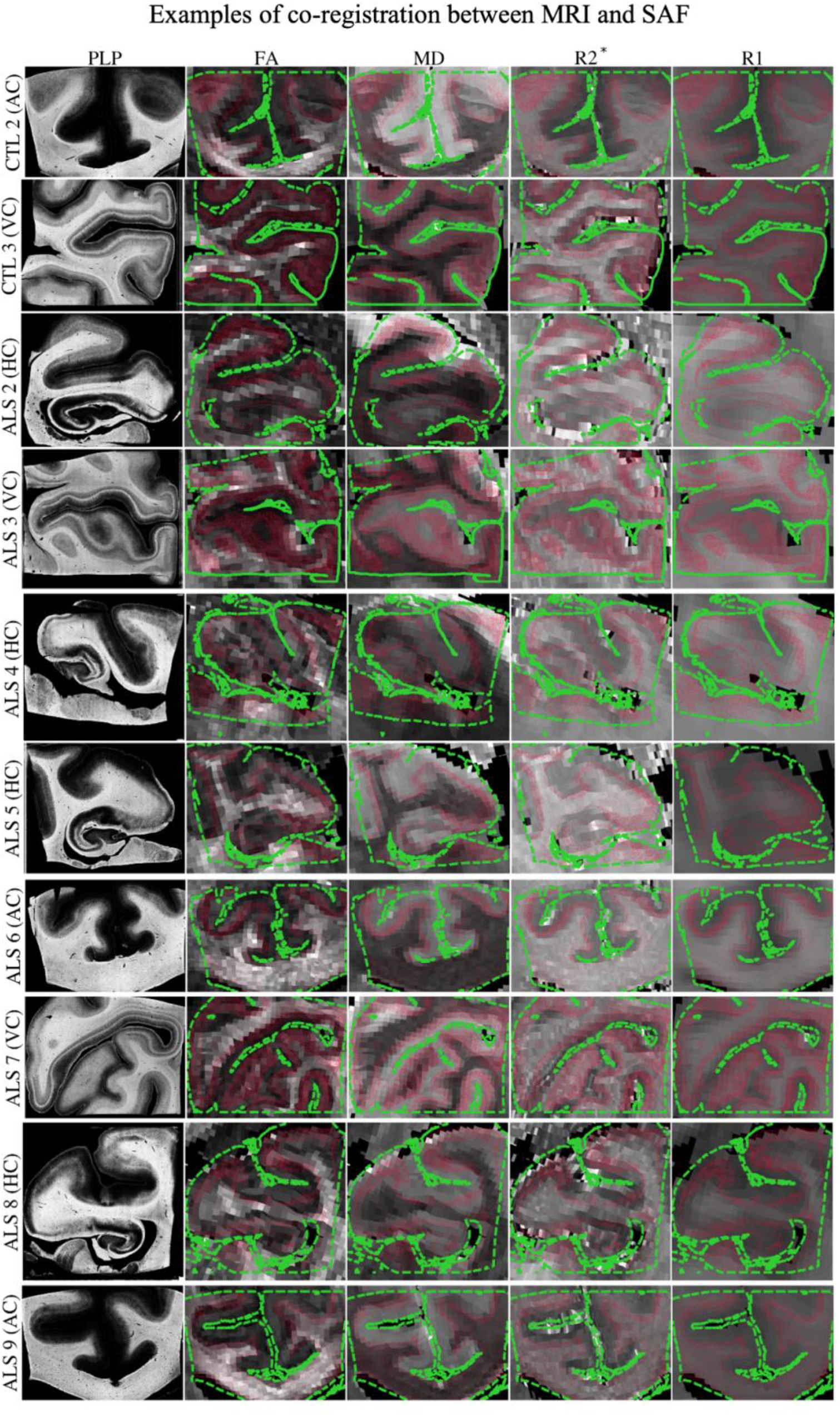
Registration evaluation for example brain regions extracted from 10 additional subjects (2 CTL, 10 ALS) (rows). In each brain region, contours of the tissue mask (green dashed) are overlaid on the co-registered MR parameter maps (second to fifth columns). The white and grey matter interface is shown in red. The tissue boundaries are closely aligned and the high registration accuracy enables us to perform meaningful voxelwise MRI-histology correlations.

## Notes

### Competing Interest Statement

The authors have declared no competing interest.

### Summary of Updates

The manuscript has been restructured to make the content easier to follow. No new results have been added.

